# Integrative Analysis of Histological Textures and Lymphocyte Infiltration in Renal Cell Carcinoma using Deep Learning

**DOI:** 10.1101/2022.08.15.503955

**Authors:** Otso Brummer, Petri Pölönen, Satu Mustjoki, Oscar Brück

**Affiliations:** Hematoscope Lab, Helsinki University Hospital, Helsinki, Finland; Hematology Research Unit Helsinki, University of Helsinki and Helsinki University Hospital Comprehensive Cancer Center, Helsinki, Finland; Translational Immunology Research Program, University of Helsinki, Helsinki, Finland; Department of Pathology, St. Jude Children’s Research Hospital, Memphis, TN, USA; iCAN Digital Precision Cancer Medicine Flagship, Helsinki, Finland; Department of Clinical Chemistry and Hematology, University of Helsinki, Helsinki, Finland

## Abstract

Evaluating tissue architecture from routine hematoxylin and eosin-stained (H&E) slides is prone to subjectivity and sampling bias. Here, we extensively annotated ∼40,000 images of five tissue texture types and ∼25,000 images of lymphocyte quantity to train deep learning models. We defined histopathological patterns in over 400 clear-cell renal cell carcinoma H&E-stained slides of The Cancer Genome Atlas (TCGA) and resolved sampling and staining differences by harmonizing textural composition. By integrating multi-omic and imaging data, we profiled their clinical, immunological, genomic, and transcriptomic phenotypes. Histological grade, stage, adaptive immunity, the epithelial-to-mesenchymal transition signature and lower mutation burden were more common in stroma-rich samples. Histological proximity between the malignant and normal renal tissues was associated with poor survival, cellular proliferation, tumor heterogeneity, and wild-type *PBRM1*. This study highlights textural characterization to standardize sampling differences, quantify lymphocyte infiltration and discover novel histopathological associations both in the intratumoral and peritumoral regions.

## INTRODUCTION

Histopathological examination of hematoxylin and eosin-stained (H&E) tissue sections remains one of the cornerstones in the diagnostics and prognostics of human cancers. In computational histopathology, image analysis algorithms are trained for detection and classification tasks generally performed by pathologists^1^. Deep learning-based models such as convolutional neural networks (CNNs) and visual transformers have been able to identify both standard histological patterns such as tumor grade^2^ and mitosis^3^ but also novel patterns related to genomic alterations^4–6^, gene expression^7,8^ and viral tumorigenesis^9^.

Clear-cell renal cell carcinoma (ccRCC) is treated with partial or radical nephrectomy, while advanced diseases are managed with vascular endothelial growth factor-targeted therapy and immune checkpoint inhibitors^10^. Given the promising efficacy of immunotherapy as adjuvant^11^ or first-line combination therapy^12^, computational histopathology could provide an inexpensive tool to identify patients associated with poor survival or treatment sensitivity such as by quantifying tumor-infiltrating lymphocytes to predict immunotherapy response^13^.

With over 10,000 diagnostic slides and associated genomic, transcriptomic, epigenomic, and clinical data from 32 cancer types, The Cancer Genome Atlas (TCGA) is a unique multi-omics archive for biomedical research^14^. Previous reports on ccRCC genetics have illustrated remodeling of cellular metabolism^15^, abundant indel mutation^16^ and adaptive immunity^17^. TCGA image data have been used to indicate that *PBRM1*-mutated ccRCC samples can be identified from H&E images^7^, and that combination of genomic and image data can improve prognostication compared to standard staging^18,19^.

Histologically, ccRCC is characterized by nests of large tumor cells with a conspicuous cytoplasm and surrounded by stromal, vascular, necrotic, adipose, and normal renal tissue and immune cells. Here, we quantified tissue textures and lymphocyte infiltration in TCGA H&E-stained ccRCC digital sections and integrated clinical, genomic, immune and transcriptomic data to study the intratumoral and peritumoral histopathological landscape.

## RESULTS

### CNNs can reliably detect tissue textures and lymphocyte proportion

We trained CNNs to detect textures and lymphocyte infiltration in H&E-stained diagnostic tissue sections of TCGA ccRCC patients (Fig. 1a-b). First, we thoroughly annotated 39,458 image tiles to one of six distinct texture classes and 25,095 tiles to low or high lymphocyte classes (Fig. 1c-d). The 18-layered ResNet (ResNet-18) correctly classified 95.0% of lymphocyte tiles in the test set (n=2,510). The ResNet-34 achieved 95.7% and the ResNet-50 95.3% classification accuracy indicating that increasing model depth slowed training but did not improve model fitness. Detailed classification metrics are visualized in Fig. 1e and Extended Data Fig. 1.

**Figure 1.**
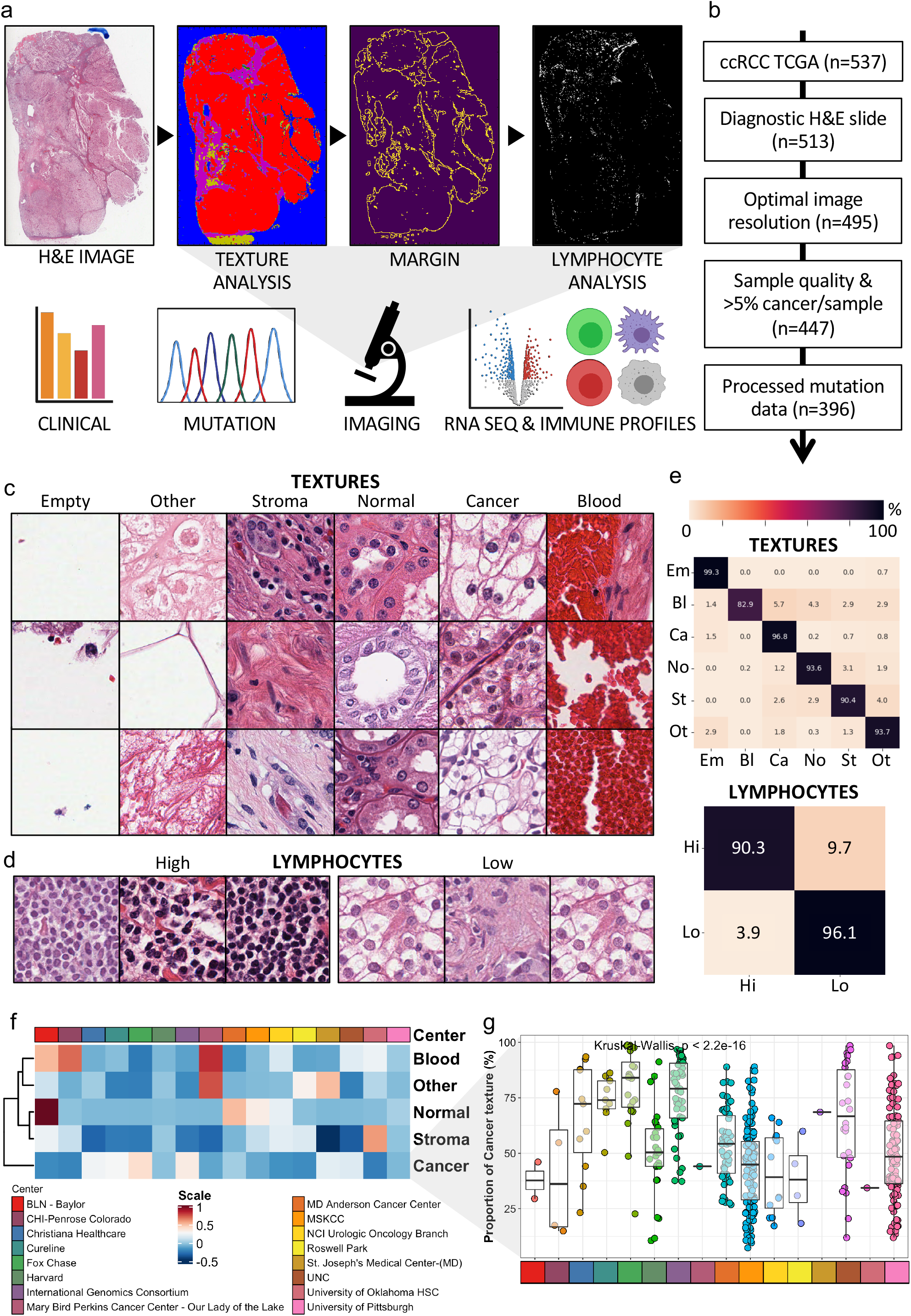
Study design. (a) Digital hematoxylin and eosin (H&E)-stained slides of clear-cell renal cell carcinoma (ccRCC) patients were collected from The Cancer Genome Atlas (TCGA). Six texture subtypes, the tumor margin and its surrounding non-margin regions as well as lymphocyte proportion were detected and quantified with convolutional neural networks. Imaging data were integrated with clinical, genomic, transcriptomic and transcriptome-based immune profiling data. (b) Flow chart of the patient number included in this study. (c) Image tile examples annotated by texture subtypes and (d) lymphocyte proportion. (e) Confusion matrix of the classification accuracy of the texture (upper) and lymphocyte (lower) classifiers. (f) Heatmap of median proportion of texture subtypes by participating TCGA clinical site. (g) Scatter and box plots to compare the proportion of cancer tissue textures by participating TCGA clinical site. The box plots indicate the interquartile ranges and median values.

For consistency, we used the ResNet-18 infrastructure also for texture classification. The algorithm achieved 95.8% total classification accuracy in the test set (n=3,946) ranging from 82.9% for blood texture to 99.3% to empty areas (Fig. 1e). Most of the classification errors were related to tiles composed of a mixture of textures. For instance, hemorrhage is a typical histopathological finding in ccRCC due to neovascularization and structural instability of rapidly growing blood vessels^25^. Therefore, the misclassification of blood tiles as cancer texture was likely due to their co-occurrence. The algorithm misclassified only 2 out of 1,212 (0.2%) cancer texture images as normal renal tissue and 8 out of 645 (1.2%) normal renal tissue images as renal cancer indicating excellent distinction.

### TCGA clinical sites differ by their texture characteristics

TCGA ccRCC samples have been collected from 16 participating clinical sites. The tissue procurement protocol has been described^14^. However, the actual sampling uniformity remains unknown, although this may significantly hamper analysis of TCGA samples.

By comparing the texture composition of samples by their clinical site, we observed that the proportion of cancer texture varied 4-fold between 22.0-80.0% (Fig. 1f-g). We noted substantial differences also in other texture types (Extended Data Fig. 2). A median of >5% normal renal tissue was evident only in samples originating from the MD_Anderson_Cancer_Center, BLN_Baylor and MSKCC indicating that histological slides have not been submitted to TCGA with the same protocol (Fig. 1f and Extended Data Fig. 2).

We excluded 51 samples from 48 patients (9.6%) as these did not represent typical ccRCC histology or included <5% cancer texture (Fig. 1b and Supplementary Table 1). The reason for low cancer proportion was atypical or non-ccRCC histology (n=22), poor histological quality (n=21), necrotic sample (n=4), and lack of cancer tissue (n=4). These findings highlight the priority of computational review to ensure high sample quality.

### Tissue hemorrhage is associated with lower metastasis rate, less frequently mutated mTOR and lower infiltration of regulatory T-cells

Next, we aimed to resolve the textural landscape of ccRCC patients and the determinants regulating its composition. As expected, renal cancer tissue was the most prevalent texture (median 49.6% IQR [33.2%-68.8%]), followed by stroma (13.0% [7.3-21.7]), other (11.6% [5.3-24.8]), blood (2.7% [0.8-7.8]) and normal tissue (2.4% [0.2-11.9]; Fig. 2a and Extended Data Fig. 3a). In total, in 227/447 (50.8%) samples the tissue section was covered by less than 50% of cancer cells implying high textural heterogeneity.

**Figure 2.**
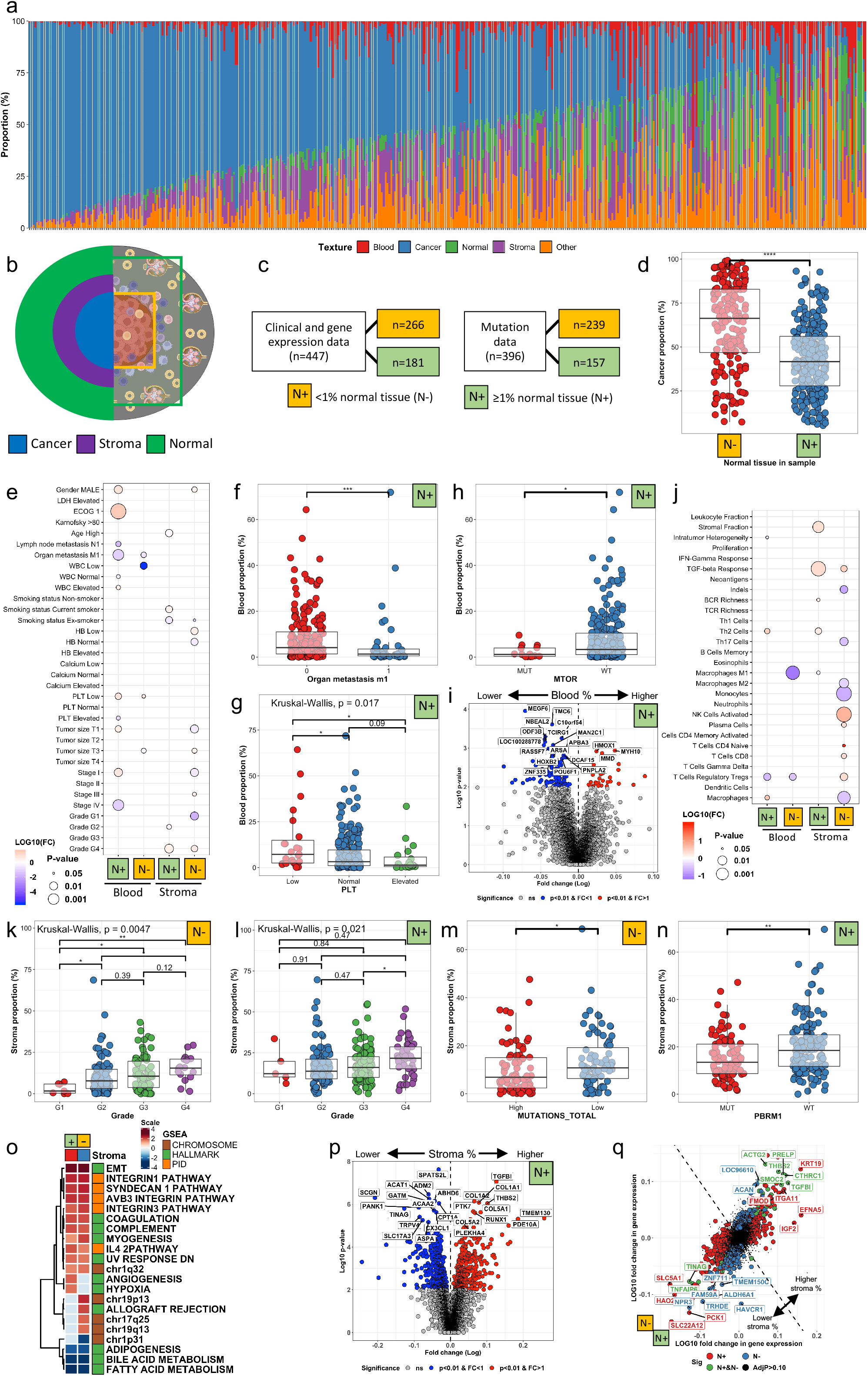
Tissue texture analysis. (a) Barplot of tissue texture profiles (y-axis) in individual patients (x-axis, n=447). (b) Schematic of the clear-cell renal cell carcinoma microenvironment. The left side illustrates three common subregions by their principal texture type: the intratumoral cancer tissue, the stroma-rich margin and the outer normal renal tissue. The right side illustrates two examples of sampling conventions: without (green rectangle) or with (light blue rectangle) normal renal tissue. (c) Flow chart of patient numbers divided by the proportion of normal tissue. (d) Scatter and box plots to compare the proportion of cancer texture by samples without or with normal tissue. (e) Balloonplot illustrating the association between clinical variables (y-axis; binary variables) and texture proportions (x-axis; continuous variables). (f) Scatter and box plots to compare the proportion of blood tissue texture by organ metastasis, (g) peripheral blood platelet level and (h) *mTOR* mutation status in samples with normal tissue. (i) Volcano plot comparing the genes differentially expressed in tumors with higher (right) or lower (left) than median red blood cell proportion. (j) Balloonplot illustrating the association between transcriptome-based signatures (y-axis; binary variables) and texture proportions (x-axis; continuous variables). (k) Scatter and box plots to compare the proportion of stroma by tumor Fuhrman grade in samples without and (l) with normal tissue. (m) Scatter and box plots to compare the proportion of stroma by mutation burden in samples without normal tissue. (n) Scatter and box plots to compare the proportion of stroma by *PBRM1* mutation status in samples with normal tissue. (o) Heatmap of the normalized enrichment score of the significant gene pathways associated (adjusted p<0.05) with samples with and without normal tissue. Grey-colored boxes indicate the pathways associated with only samples with or without normal tissue (adjusted p<0.05) but not with the other (adjusted p>0.10). Only Chromosome, Hallmark and PID pathways have been included in the analysis to simplify visualization. (p) Volcano plot comparing the genes differentially expressed in tumors with higher (right) to lower (left) than median stroma proportion. Only samples with normal tissue are included. (q) Scatter plot showing the preferential association with higher stroma tissue proportion in samples with or without normal tissue. Box plots: the lower and upper limit indicate the interquartile range and the center line the median value. Balloonplots: Only significant associations are marked with balloons. The color of the balloon reflects the association power (log10 fold change) and its size the statistical significance. LOG10(FC); 10-fold logarithmic value of the texture fold change by clinical markers for example gender (ratio of the median blood proportion in male:female).

Besides tumor biology, the varying site-specific sampling protocols likely affected the textural content. We reasoned that by dividing TCGA ccRCC samples into those without (<1%) and with normal tissue (≥1%) we could examine tissue textures in two distinct but histologically more coherent cohorts (Fig. 2b-c). Samples without normal renal tissue (N-) were composed of 66.3% [IQR 46.9-82.8%] cancer texture representing the tumor core compared with 41.6% [27.9-56.0%] in samples with normal renal tissue (N+) reflecting a broader tumor microenvironment (Fig. 2d and Extended Data Fig. 3b-d).

As the proportion of normal renal tissue reflected tissue sampling practices and the “other” texture class included various histological types, we focused on blood and stroma-associated phenotypes. Higher hemorrhage in N+ samples was associated with lower organ metastasis, lower tumor stage and superior Eastern Cooperative Oncology Group (ECOG) performance status (Fig. 2e-f). We also noted that the peripheral blood (PB) platelet count gradually decreased with increasing proportion of tumor hemorrhage in N+ samples (Fig. 2g). Elevated pretreatment platelet level is a biomarker of poor survival and is incorporated in the prognostic Heng score^26^. Thus, low PB platelet count could be due to high angiogenic activity and consequent tumor hemorrhage.

When examining N- samples, we observed similar but less significant association of tumor hemorrhage and lower rate of organ metastasis and lower PB platelet count (Fig. 2e). Tumor hemorrhage was not associated with overall survival in either N+ or N- cohorts (Extended Data Fig. 4).

We investigated next genomic alterations. In N+ samples, *mTOR*^mut^ occurred in 24/396 patients (6.1%) and associated with lower hemorrhage (Fig. 2h). While mTOR has been described to increase angiogenesis via HIF1α and VEGF-regulated pathways, *VHL*^mut^ was not associated with hemorrhage indicating another mechanism. No association with other gene alterations, mutation burden or aneuploidy was observed (Supplementary Table 2). No association between tumor hemorrhage and genomic variables were observed in N- samples (Supplementary Table 3).

To identify hemorrhage-associated transcriptomic signatures, we first compared the expression of individual genes (Fig. 2i and Extended Data Fig. 5a). In N+ samples, the *HMOX1* gene was highly expressed in conjunction with hemorrhage (Fig. 2i). *HMOX1* encodes the heme oxygenase 1 enzyme catalyzing heme to biliverdin and increased heme catabolism is consistent with increased tissue hemorrhage^27^. We observed upregulated inflammatory, epithelial-to-mesenchymal (EMT) and hypoxia-related pathways in samples without normal tissue but little difference in immune profiles (Fig. 2j and Extended Data Fig. 5b-c). In summary, our findings indicate that peritumoral and intratumoral hemorrhage are histologically distinct phenotypes by their clinical, mutational, transcriptional, and immunological profiles.

### Tissue fibrosis is associated with high histological grade, low mutation burden and an adaptive immune response

Next, we examined fibrosis-related manifestations. In N- samples, we observed association between stroma and histological grade (Fig. 2e and Fig. 2k). Similar relation was less evident in N+ samples indicating that intratumoral but not peritumoral stroma would be linked with poor renal cell differentiation (Fig. 2l). In line, N- stroma associated with other established adverse prognostic biomarkers such as tumor size, stage and anemia (Fig. 2e).

When studying genomic alterations, mutation burden was linked with lower histological fibrosis both in N+ and N- samples (Fig. 2m and Supplementary tables 4-5). In N+ samples, *PBRM1*^wt^ and diploid haplotype were also associated with higher proportion of stroma (Fig. 2n and Supplementary table 4). In N- samples, *SETD2*^mut^ indicated lower fibrosis (Supplementary table 5).

Fibrosis in N- samples was associated with an adaptive immune signature characterized with enriched CD8+ T-cells, regulatory T-cells and activated NK-cells (Fig. 2j and Extended Data Fig. 5d-f). Moreover, the proportion of M2-polarized macrophages and Th17-differentiated helper T-cells were reduced (Fig. 2j and Extended Data Fig. 5g-i). As mutation burden and *PBRM1* genotype have been implicated with immunotherapy response, quantifying fibrosis could provide an inexpensive biomarker to increase their precision or select patients for targeted sequencing^28–30^.

Next, we inspected fibrosis-associated transcriptional programs. As expected, the TGF β response pathway was enriched both in N+ and N- samples with high stromal composition (Fig. 2j). High histological stroma associated only in N+ samples with transcriptome-derived stromal score (Fig. 2j). We reasoned this to be due to more abundant stroma in N+ compared to N- samples as visually confirmed by large peritumoral stromal margins compared to intratumoral fibrotic islets (Extended Data Fig. 6a).

Established stromal signaling pathways regulating EMT and the formation of integrin, syndecan, and collagen were elevated in both N+ and N- samples (Fig. 2o). Moreover, fibrosis was associated with coagulation and less active lipid metabolism (Fig. 2o). The chromosomal locus 19p13 was overactivated in conjunction with enriched stroma in N- samples, while hypoxia and angiogenesis-related pathways were elevated in N+ samples with high stroma (Fig. 2o). Similar findings were observed also at the gene-level (Fig. 2p and Extended Data Fig. 6b). While many of genes were associated with stroma formation in both sample categories, integrin alpha 11 (*ITGA11*), ephrin 5 (*EFNA5*) and cytokeratin 19 (*KRT19*) were enriched in N+ samples whereas aggrecan (*ACAN*) in N- samples. These findings suggest that the extracellular matrix (ECM) in the intratumoral and peritumoral tissue could differ by their adhesion, migration and cell signaling abilities (Fig. 2q).

### Lymphocyte infiltration is coordinated between malignant and surrounding tissue

Previous studies quantifying tumor lymphocyte infiltration in ccRCC have relied on deconvolution of bulk RNA-sequencing data^17^, histochemical,^31^ or antibody-based detection such as flow cytometry^32,33^. While these approaches are precise to approximate the lymphocyte population in a sample, they impose demands on uniform sampling and sample processing.

Here, we built a deep learning model identifying images with high (90.3% classification accuracy) and low (96.1%) lymphocyte density in the test set (Fig. 1e). We hypothesized that texture-aware lymphocyte quantification could solve issues related to sampling variation. The lymphocyte classification probability per tile reflected lymphocyte density. Therefore, lymphocyte predictions [0-1] were proportioned by texture surface area. The median lymphocyte proportion per sample was 20.3% and varied between 2.3-82.5%. The highest lymphocyte density was unexpectedly in the normal renal texture followed by cancer, stroma, blood, and lastly other texture types (Fig. 3a). Texture-specific lymphocyte proportions shared high positive correlation (Fig. 3b). Intratumoral infiltration explained most of the total sample infiltration variance (0.85^2^ = 73%; Fig. 3b).

**Figure 3.**
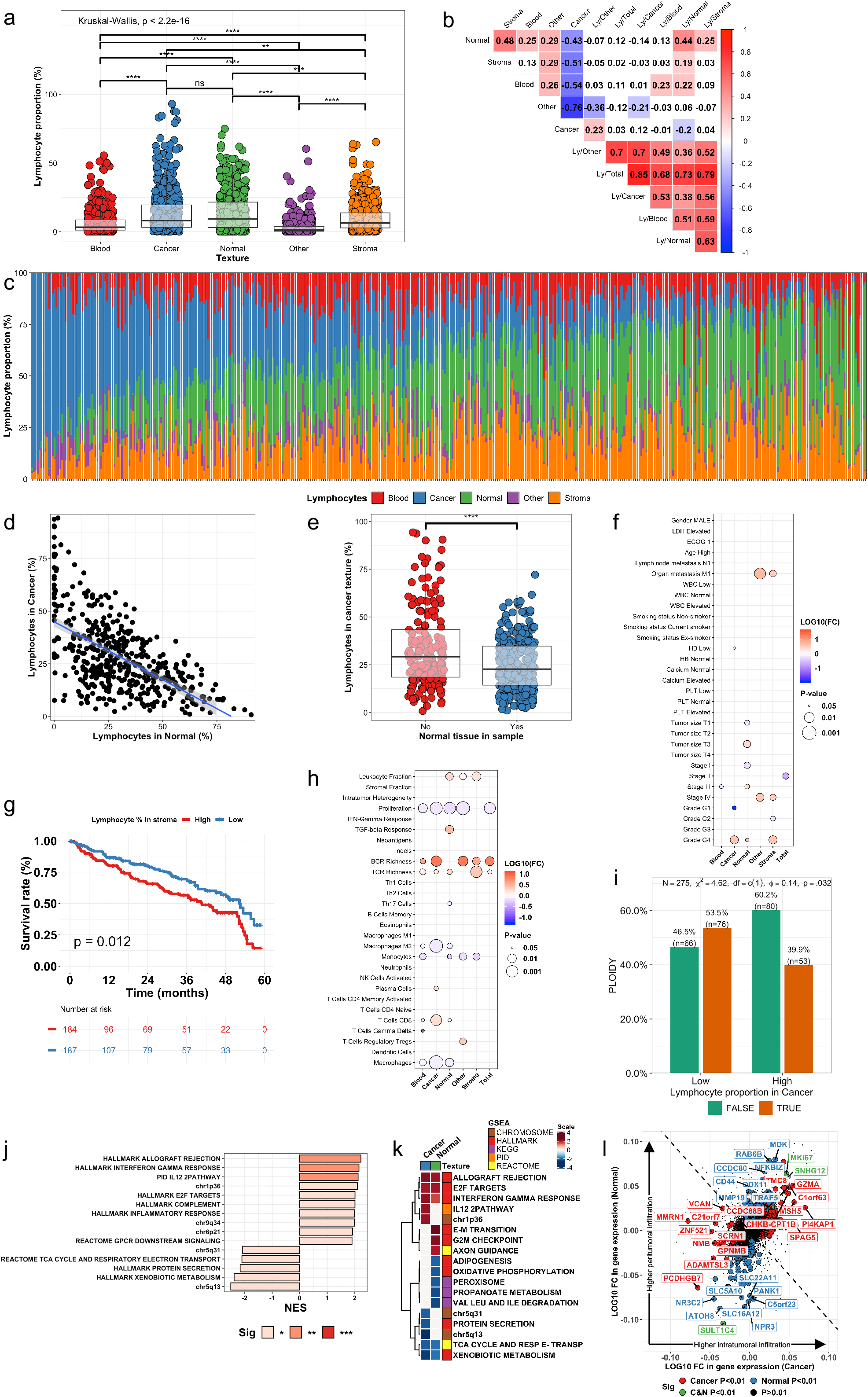
Texture-specific lymphocyte analysis. (a) Scatter and box plots to compare the proportion of lymphocytes by textures. (b) Correlation matrix of the texture and texture-specific lymphocyte proportion. (c) Barplot of relative lymphocyte proportion (density normalized to 100%) by textures (y-axis) in individual patients (x-axis, n=447). The lymphocyte density (proportion/area) has been median-averaged by texture type and then their sum rescaled to 100%. (d) Scatter plot to illustrate the correlation of relative lymphocyte proportion in malignant and normal renal tissue. (e) Scatter and box plots to compare the proportion of lymphocytes in cancer tissue by samples without or with normal tissue. (f) Balloonplot indicating the association between clinical variables (y-axis; binary variables) and texture-aware lymphocyte proportions (x-axis; continuous variables). (g) Kaplan-Meier curves showing the survival association by the proportion of lymphocytes in stroma tissue. (h) Balloonplot indicating the association between transcriptome-based signatures (y-axis; binary variables) and texture-specific lymphocyte proportions (x-axis; continuous variables). (i) Barplot illustrating the ploidy status (FALSE = aneuploidy, TRUE = normal) by lymphocyte infiltration in cancer tissue. (j) Barplot indicating the normalized enrichment score (NES) of the gene pathways significantly associated with cancer lymphocyte proportion. (k) Heatmap of the normalized enrichment score of the significant gene pathways associated (adjusted p<0.05) with lymphocyte infiltration to cancer or normal textures. Grey-colored boxes indicate the pathways associated with only samples with or without normal tissue (adjusted p<0.05) but not with the other (adjusted p>0.10). (l) Scatter plot of genes associated with lymphocyte infiltration to the cancer (red) or normal (blue) or both (green) texture. The coordinate of each dot represents the ratio of the gene expression value when comparing in (cancer or normal) texture-specific lymphocyte-rich vs. lymphocyte-low samples. Box plots: the lower and upper limit indicate the interquartile range and the center line the median value. Balloonplots: Only significant associations are marked with balloons. The color of the balloon reflects the association power (log10 fold change) and its size the statistical significance. LOG10(FC); 10-fold logarithmic value of the lymphocyte fold change difference by clinical markers for example gender (ratio of the median blood proportion in male:female).

Next, we rescaled the texture-specific lymphocyte densities so that their sum would equal to 100% and observed heterogenous distribution at the patient-level (Fig. 3c). Strikingly, the intratumoral lymphocyte density correlated negatively with the density in the surrounding normal renal tissue (R -0.56, p<0.001; Fig. 3d) and stromal texture (R-0.37, p<0.001) indicating controlled movement between the tumor and its surrounding textures.

### Stromal lymphocytes are associated with poor survival and high T-cell receptor diversity

Texture-specific lymphocyte proportions differed by the proportion of normal renal tissue (<1% vs. ≥1%; Fig. 3e and Extended Data Fig. 7a-c). To equalize sampling differences, we normalized lymphocyte density with texture-specific weights (see Methods) successfully reducing differences in the total lymphocyte infiltration by clinical center (Extended Data Fig. 7d-e). However, samples originating from Fox Chase contained higher lymphocyte density than expected (Extended Data Fig. 7e). When examined visually, these samples were characterized with a high hematoxylin:eosin ratio and lymphocyte scoring even visually was ambiguous (Extended Data Fig. 7f-g). As a conclusion, these samples were excluded from lymphocyte analyses.

To fully normalize staining differences, we categorized samples into “High” and “Low” infiltration based on their lymphocyte density compared to the clinical center median density. When examining clinical significance, we observed that lymphocyte infiltration in cancer and stromal textures was associated with high histological grade (Fig. 3f). Only stromal lymphocyte infiltration was related with poor overall survival, organ metastasis and stage IV disease (Fig. 3f-g and Extended Data Fig. 8). Patient gender, age, smoking status, or laboratory values were not related to lymphocyte infiltration (Fig. 3f).

We then studied genomic and immunological profiles. In line with our expectations, infiltration in malignant renal tissue was associated with the transcriptomic CD8+ T-cell signature (Fig. 3h). We also observed a CD8+^high^/M2-macrophage^low^ combination in patients with high lymphocyte infiltration notably in the cancer texture (Fig. 3h). B and T-cell receptor diversity was associated with increased lymphocyte infiltration almost irrespective of textural context (Fig. 3h). Of note, cell proliferation correlated negatively with all except stromal lymphocyte infiltration (Fig. 3h).

### Aneuploidy, chromosome 1p and 5q loci, and the EMT program identified as regulators of tumor-infiltrating lymphocytes

Next, we examined genomic alterations associated with lymphocyte infiltration. Aneuploidy was associated with higher intratumoral and total lymphocyte density (Fig. 3i and Extended Data Fig. 9a). Mutation burden was related to higher infiltration to blood texture (Extended Data Fig. 9b). When studying individual genes, *SETD2*^mut^ trended with higher infiltration in stromal, other and blood textures (Extended Data Fig. 9c-e). The *PBRM1* genotype has been associated with both nonimmunogenic^30^ and immune hot phenotype and anti-PD1 therapy response^29^. In our study, *PBRM1* alterations were not associated with total or texture-specific lymphocyte infiltration (Extended Data Fig. 9f-k). By analyzing the supplementary data provided by Braun *et al*^29^, no association was evident between the tumor core CD8+ density and *PBRM1* status validating our finding and indicating that its immunologic significance remains unclear (Extended Data Fig. 9l).

Finally, we studied the transcriptomic programs associated with lymphocyte infiltration. The top three pathways enriched with intratumoral infiltration were well-established T-cell activation signatures endowing confidence to our analysis (Fig. 3j). The chromosomal locus 1p36 was the next most overactivated pathway while the 5q31 and 5q13 loci and metabolic pathways were most downregulated (Fig. 3j).

Given our previously described asynchronous lymphocyte enrichment in either malignant or normal renal tissue, we interrogated which pathways are most commonly altered in these two compartments. Adaptive immune response pathways were activated in lymphocyte-rich samples (Fig. 3k). Exclusively in samples with lymphocyte-rich cancer texture, the IL-12 pathway, and genes of the chromosome 1p36 locus were upregulated and genes of the chromosome 5q13 and 5q31 downregulated. On the contrary, the hallmark EMT pathway was enriched, while lipid and energy production pathways depleted in lymphocyte-rich normal renal tissue (Fig. 3k). The cytolytic granzyme A (*GZMA*) enzyme illustrious of T and NK-cells and the testis antigen *SPAG5* were associated with infiltration to cancer tissue (Fig. 3l). On the contrary, genes regulating the ECM and fibrosis such as *MMP19* and *CD44* were upregulated in samples with infiltration to normal renal tissue (Fig. 3l). In summary, immune, mesenchymal, and metabolic factors influence lymphocyte infiltration.

### The textural composition of the tumor margin predicts prognosis and reflects the tumoral genomic and transcriptomic alterations

Based on our previous findings that the tumor core and its surrounding peritumoral tissue form two immunologically distinct regions^34^. Therefore, we were intrigued to investigate the textural content of the peritumoral margin and its exterior non-margin region (Fig. 4a).

**Figure 4.**
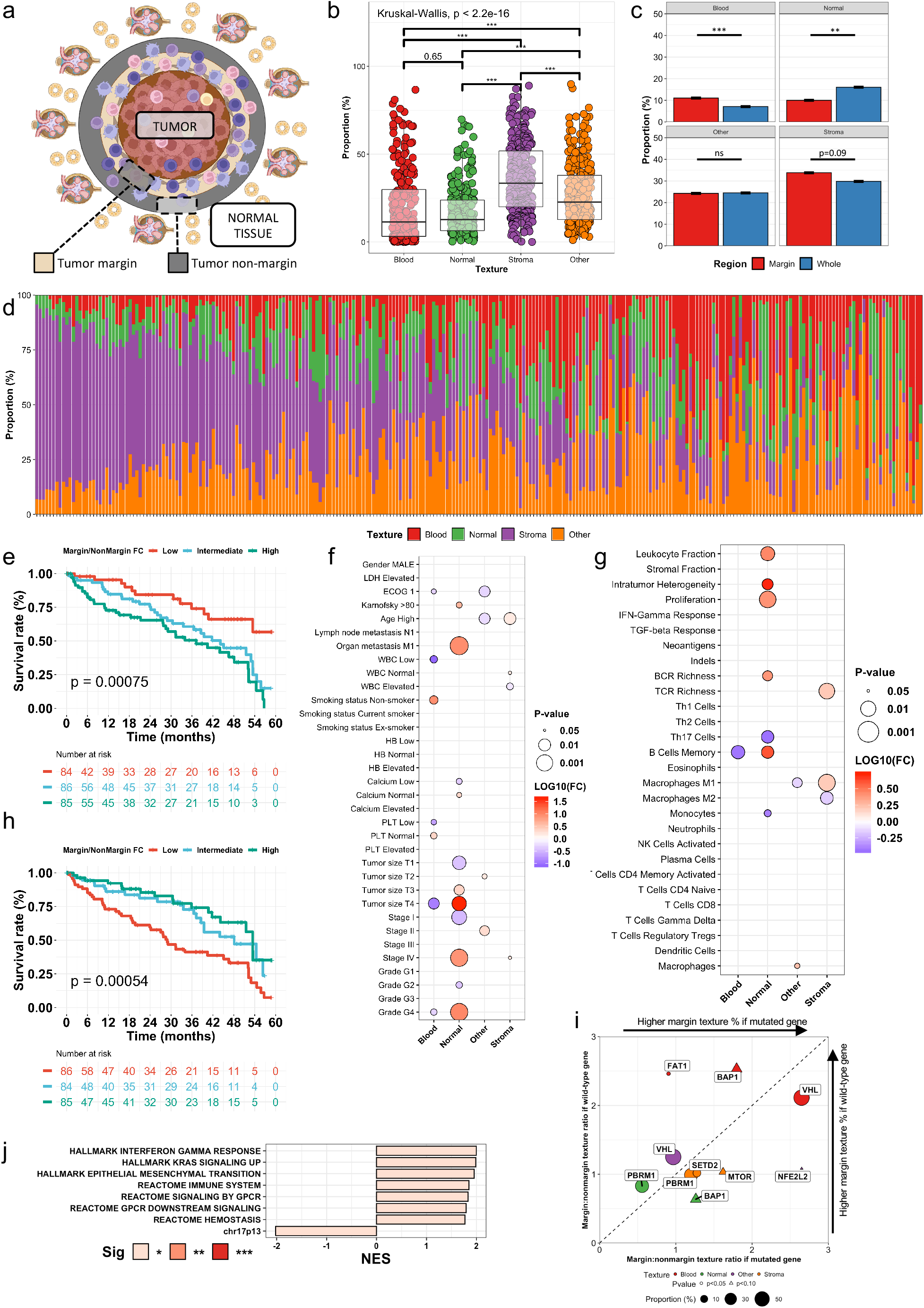
Texture profile in the tumor margin. (a) Schematic of the peritumoral tissue consisting of an inner margin and outer non-margin tissue. (b) Scatter and box plots to compare the proportion of tissue textures in the tumor margin. (c) Box plots to compare the proportion of tissue textures in the tumor margin and the entire sample. (d) Barplot of the relative texture proportion (y-axis) in the tumor margin in individual patients (x-axis, n=447). (e) Kaplan-Meier curves showing the survival association by the margin:non-margin ratio of the proportion of normal tissue. (f) Balloonplot indicating the association between clinical variables (y-axis; binary variables) and the margin:non-margin ratio of four textures (x-axis; continuous variables). (g) Balloonplot indicating the association between transcriptome-based signatures (y-axis; binary variables) and the margin:non-margin ratio of four textures (x-axis; continuous variables). (h) Kaplan-Meier curves showing the survival association by the margin:non-margin ratio of the proportion of blood tissue. (i) Scatter plots illustrating enrichment of genomic alterations by margin:non-margin ratio of tissue proportions. Only associations with p<0.10 are visualized. (j) Barplot indicating the normalized enrichment score (NES) of the gene pathways significantly associated with the ratio of margin:non-margin stroma proportion. Box plots: the lower and upper limit indicate the interquartile range and the center line the median value. Balloonplots: Only significant associations are marked with balloons. The color of the balloon reflects the association power (log10 fold change) and its size the statistical significance. LOG10(FC); 10-fold logarithmic value of the texture fold change difference by clinical markers for example gender (ratio of the median blood proportion in male:female).

The peritumoral texture was dominated by stroma and followed by textures in the same order as observed in the entire sample (Fig. 4b). Blood and stroma textures were more frequent in the tumor margin compared to both the non-margin region and the entire sample (Fig. 4c and Extended Data Fig. 10a). In contrary, normal renal tissue was less common in the tumor margin than in the entire sample (Fig. 4c and Extended Data Fig. 10a). These findings are concordant with previous findings on the fibrous capsule and tumor-penetrating stromal islets (Extended Data Fig. 6).

The margin composition was heterogenous at the patient-level (Fig. 4d). We assigned each patient with a textural enrichment score by comparing texture proportions between the tumor margin to the non-margin tissue. This approach was unaffected of technical differences in tissue staining as two regions of the same sample are compared. Patients with a high margin:nonmargin normal renal tissue ratio had significantly worse survival (Fig. 4e). The clinical profile of these patients included higher tumor size, histological grade, organ metastasis and advanced stage (Fig. 4f). These tumors shared also higher proliferation, heterogeneity and leukocyte fraction including more frequent and clonally diverse memory B-cells (Fig. 4g).

In the opposite, patients with elevated blood texture in the tumor margin were characterized with superior survival (Fig. 4h). These patients shared known favorable clinical characteristics such as lower tumor size, more differentiated tumor cells and less frequent thrombocytopenia (Fig. 4f).

The prognosis of patients with a high peritumoral margin:nonmargin stroma did not differ from other patients, but were older at diagnosis (Fig. 4f and Extended Data Fig. 10b). Immunologically, these were characterized with a broader T-cell clonality spectrum and more frequent M1-polarized macrophages (Fig. 4g).

Next, we examined the mutational landscape of the margin histological subtypes (Fig. 4i). The margin:nonmargin normal renal tissue ratio was lower with *PBRM1*^mut^ (ratio 0.56 vs. 0.83, p=0.046) and higher with *BAP1*^mut^ (1.3 vs. 0.64, p=0.062). The mutational profile of stroma-rich margin included *PBRM1*^mut^ (1.2 vs. 0.99, p=0.0055), *SETD2*^mut^ (1.3 vs. 1.0, p=0.026) and *MTOR*^mut^ (1.6 vs. 1.0, p=0.065). We observed decreased margin tissue hemorrhage in patients with *FAT1*^mut^ (0.90 vs. 2.5, p=0.046) and *BAP1*^mut^ (1.8 vs. 2.5, p=0.082), and the opposite in patients with *VHL*^mut^ (2.6 vs. 2.1, p=0.035). Mutation burden and aneuploidy were not associated with margin texture content.

We then asked how the expression of individual genes and signaling pathways would impact the peritumoral textural composition. While the findings at single gene level were challenging to interpret, we observed distinct pathway signatures (Extended Data Fig. 10c-e). In samples with fibrotic margin, IFNγ signaling, immune responses and the EMT were activated resembling the previously described normal renal tissue lymphocyte signature (Fig. 4j). High margin:nonmargin normal renal tissue was associated with enriched cell cycle genes and downregulation of lipid and carbohydrate metabolism (Extended Data Fig. 10d). Margin:nonmargin hemorrhage correlated with downregulation of the cell cycle, chromosome 17q21, 17q25 and 6p21 loci, and MTOR and MYC signaling ascertaining the link with prognosis (Extended Data Fig. 10e).

### The lymphocyte-rich stromal margin associates with an adaptive immune response, dampened EMT and non-smoking habit

To conclude, we quantified the margin:non-margin lymphocyte ratio. The highest lymphocyte density was in the normal renal tissue (Fig. 5a). The texture order was similar than in the non-margin and the entire sample, but the lymphocyte density was higher in the tumor margin (Fig. 5b and Extended Data Fig. 11a).

**Figure 5.**
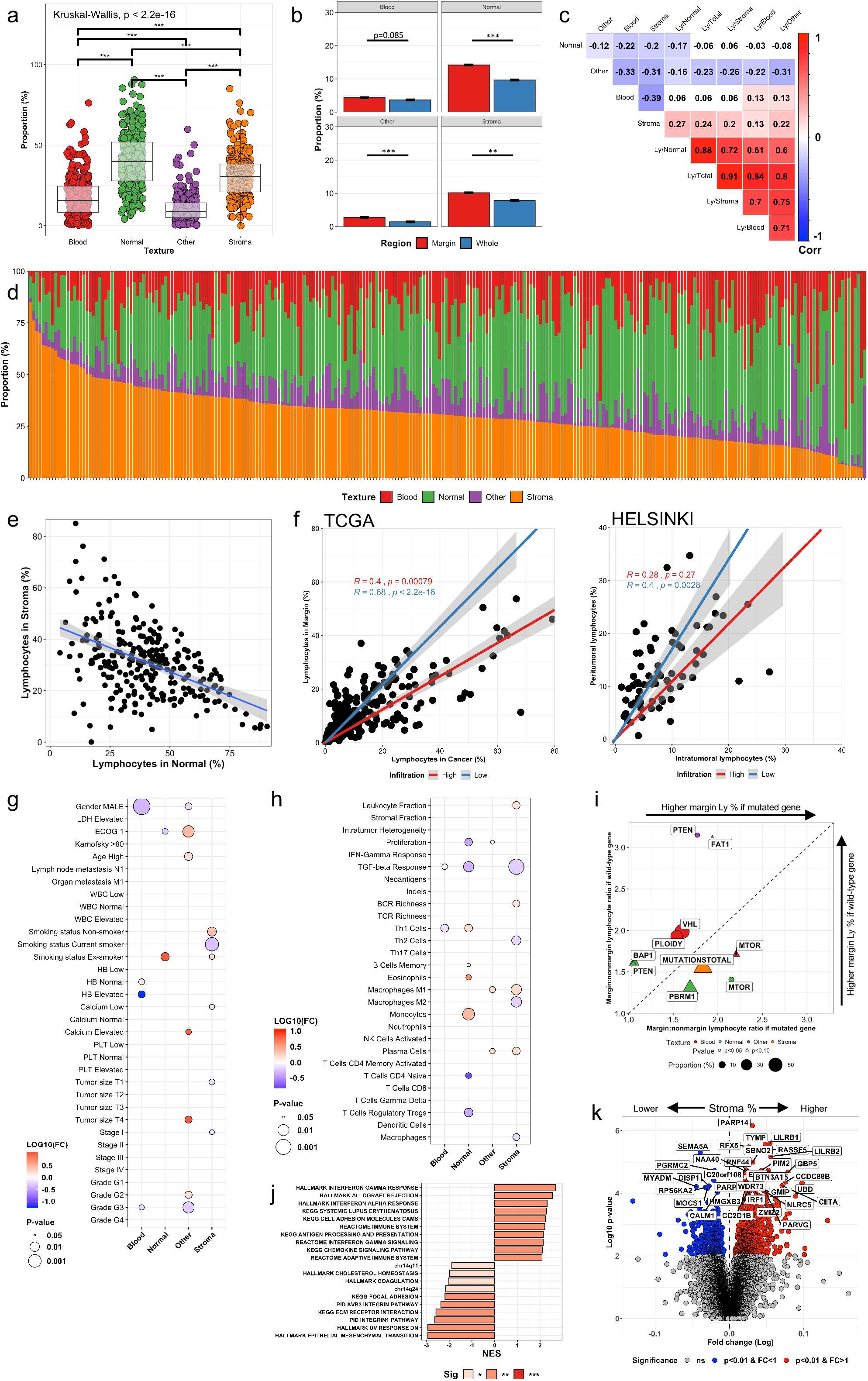
Lymphocyte infiltration in the tumor margin. (a) Scatter and box plots to compare the proportion of lymphocytes in the tumor margin by textures. (b) Box plots to compare the proportion of lymphocytes by textures in the tumor margin and the entire sample. (c) Correlation matrix of the texture and texture-specific lymphocyte proportion in the tumor margin. (d) Barplot of relative lymphocyte proportion (density normalized to 100%) by textures (y-axis) in the tumor margin of individual patients (x-axis, n=447). The lymphocyte density (proportion/area) has been median-averaged by texture type and then their sum rescaled to 100%. (e) Scatter plot and linear regression curve of the relation of relative lymphocyte proportion in stromal margin and normal renal margin. (f) Scatter plots and Spearman correlation line between the lymphocyte density in the cancer texture and tumor margin of this study and in the intratumoral and peritumoral regions of our previously published Helsinki study (Brück et al. Modern Pathology 2021). Patients are divided by highest 25% (red) and lowest 75% (blue) cancer texture lymphocyte density. (g) Balloonplot indicating the association between clinical variables (y-axis; binary variables) and the margin:non-margin ratio of lymphocyte proportion in four textures (x-axis; continuous variables). (h) Balloonplot indicating the association between transcriptome-based signatures (y-axis; binary variables) and the margin:non-margin ratio of lymphocyte proportion in four textures (x-axis; continuous variables). (i) Scatter plots illustrating enrichment of genomic alterations by margin:non-margin ratio of lymphocyte proportions in tissue textures. Only associations with p<0.10 are visualized. (j) Barplot indicating the normalized enrichment score (NES) of the gene pathways significantly associated with the ratio of margin:non-margin stroma proportion. (k) Volcano plot comparing the genes differentially-expressed in tumors with higher (right) or lower (left) margin lymphocyte density compared to non-margin lymphocyte density. Box plots: the lower and upper limit indicate the interquartile range and the center line the median value. Balloonplots: Only significant associations are marked with balloons. The color of the balloon reflects the association power (log10 fold change) and its size the statistical significance. LOG10(FC); 10-fold logarithmic value of the texture fold change difference by clinical markers for example gender (ratio of the median blood proportion in male:female).

The lymphocyte density correlated over margin textures (Fig. 5c). The stromal lymphocyte density explained most (82.8%) of the variance of total margin lymphocyte density (Fig. 5c). Elevated stroma in the margin was associated with higher margin lymphocyte density (Fig. 5c). On the contrary, the margin histological textures correlated negatively with one another indicating that if one texture type increased, the remaining textures decreased accordingly (Fig. 5c-d). This was especially evident between stroma and normal renal tissues (Fig. 5d-e).

At first, we observed high concordance between the cancer texture and peritumoral lymphocyte density (R=0.78 p<0.001, Fig. 5f). We replicated the finding in our previously reported Helsinki dataset of 64 ccRCC patients^34^ where instead of a cancer texture and its margin we had annotated the intratumoral and peritumoral regions (R=0.60 p<0.001, Fig. 5f). However, in closer inspection we could discern two linearities and, therefore, divided patients into two groups: 25% highest and 75% lowest tumoral lymphocyte density. Indeed, patients with high intratumoral infiltration had lower peritumoral lymphocyte density suggesting distinct regulatory mechanisms for lymphocyte penetration (Fig. 5f).

We studied next clinical correlates. The margin lymphocyte ratio was not clearly associated with overall survival or prognostic variables (Fig. 5g and Extended Data Fig. 11b). Infiltration in the stromal margin was decreased in active smokers and partly normalized in ex-smokers (Fig. 5g). It was also associated with a weaker TGFβ response and less abundant Th2-differentiated T-cells and macrophages (Fig. 5h). Lymphocyte abundance in the normal renal margin tissue was instead associated with monocytes and Th1 T-cells and suppression of regulatory T-cells and TGFβ pathway (Fig. 5h).

We observed that mutation burden was associated with infiltration to the stromal margin compared to the non-margin tissue (ratio 1.8 vs. 1.6, p=0.068; Fig. 5i). In addition, *VHL*^mut^ (1.6 vs. 2.0, p=0.021) and diploidy (1.5 vs. 1.9, p=0.025) were associated with lower infiltration, whereas *MTOR*^mut^ with higher infiltration to the blood-rich margin (2.2 vs. 1.7, p=0.067). Lymphocyte infiltration to the normal renal margin was associated with *MTOR*^mut^ (2.2 vs. 1.4, p=0.024), *PBRM1*^mut^ (1.7 vs. 1.3, p=0.056), *BAP1*^*wt*^ (1.6 vs. 1.1, p=0.058) and *PTEN*^*wt*^ (1.6 vs. 1.1, p=0.074).

Lastly, we studied the transcriptomic signature of the peritumoral lymphocyte infiltration (Fig. 5j, Extended Data Fig. 12). We observed activation of interferon signaling and adaptive immune response and downregulation of EMT and other mesenchymal pathways in samples with a lymphocyte-rich stromal margin (Fig. 5j). The immune-related genes included the transcriptional activator of interferon-inducible genes *IRF1*, the class II human leukocyte antigen *CIITA* required for antigen presentation and immunoregulatory genes *LILRB1* and *LILRB2* (Fig. 5k).

## DISCUSSION

The main findings of this study are 1) the texture-based approach to dissect digital histological slides and 2) the integrative network between tissue textures, lymphocyte infiltration, clinical variables, genomic alterations, and transcriptomic signatures.

TCGA tissue procurement has been described^14^, but limitedly evaluated^35^. Here, we indicate that some centers have systematically included samples spanning from the tumor core to the surrounding healthy tissue while others have restricted sampling to the tumor core. Moreover, the hematoxylin:eosin ratio differed substantially by clinical centers as previously reported^35^. While sampling differences are apparent in TCGA histological slides, these likely extend to sequencing data urging for uniform sampling, sample preprocessing, and retrospective evaluation standards.

To address the staining and sampling variance, we employed several measures (Fig. 6). The systematic dissection of ccRCC textures and lymphocyte infiltration uncovered a delicate network with connections to clinical, immunological, genomic, and transcriptomic features. For instance, while the histological grade is based on ccRCC cell morphology^36,37^, it correlated substantially with intratumoral fibrosis, which could facilitate routine tumor grading. Fibrosis correlated with immune, hypoxia and EMT signaling and alterations in commonly mutated genes (Fig. 6).

**Figure 6.**
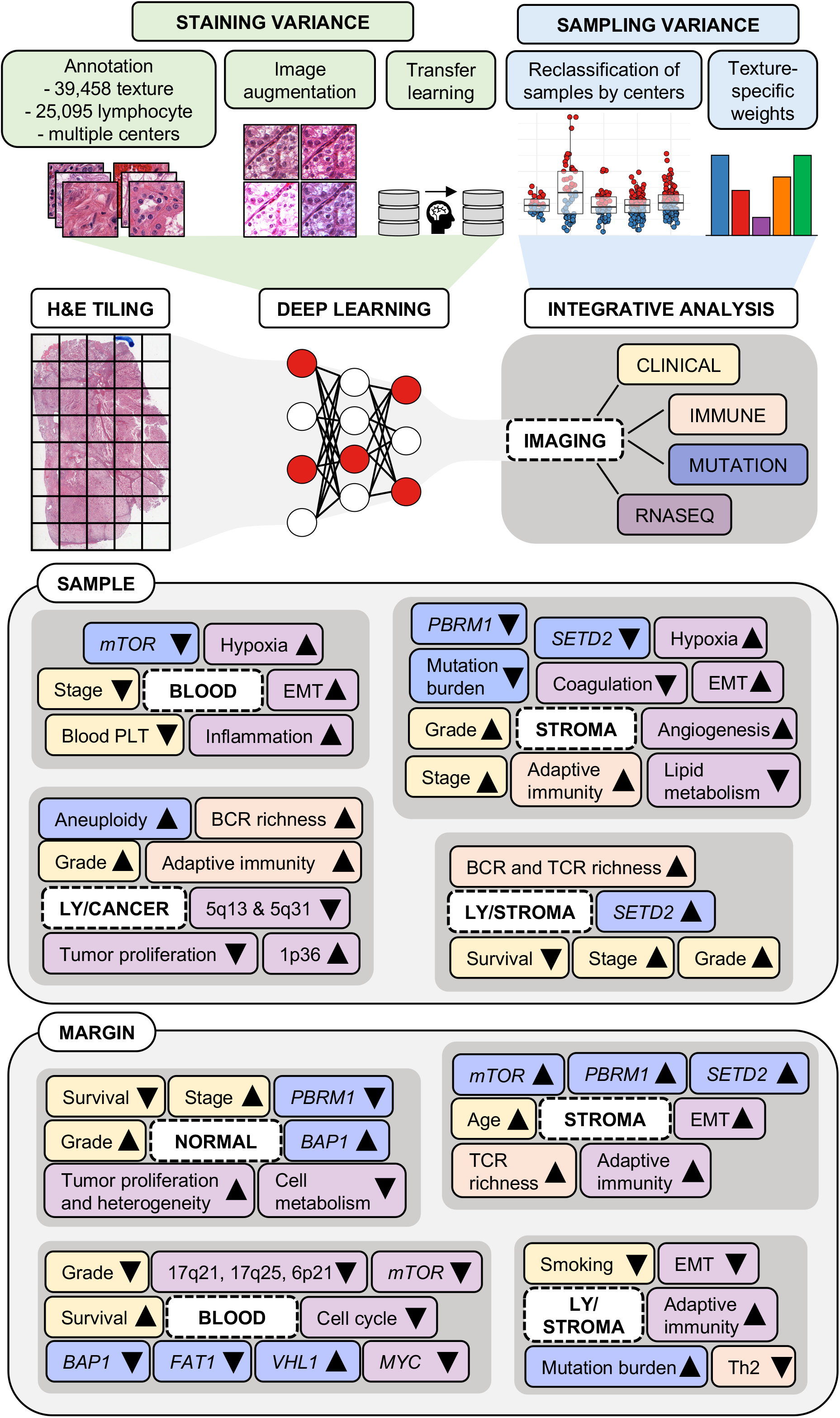
Summary of the study framework (upper) and findings (lower). Abbreviations: H&E, Hematoxylin and eosin; EMT, epithelial-to-mesenchymal transition; PLT, platelet; BCR, B-cell receptor; TCR, T-cell receptor; Ly, Lymphocyte density; Th2; type 2 helper T-cell.

Half of the lymphocytes in sections were located among cancer cells and shared moderate concordance across textures. Aneuploidy, the IL-12 pathway, chromosome 1p36 activity and chromosome 5q13 and 5q31 inactivity were linked to intratumoral infiltration. Copy number losses of 1p36 and gain of 5q35 are highly common in ccRCC, and the latter has been suggested to induce activation of the activation of PI(3)K pathway^15^. These loci might have now another immunoregulatory role.

The relative proportion of lymphocytes in cancer tissue correlated negatively with their relative proportion in the surrounding normal and stromal tissue textures possibly indicating two gravitation centers. We have previously shown that intratumoral T-cells are immunophenotypically more experienced based on higher expression of cytolytic, immune checkpoint and senescence markers and closer intercellular proximity compared to the peritumoral and normal renal regions^34^. In both the previous and current study, we demonstrated that peritumoral and intratumoral lymphocyte densities correlated if the intratumoral density was low. Possibly due to restricted lymphocyte penetration, elevated stromal lymphocyte infiltration associated with higher metastatic rate, poor survival, T and B clonotype diversity, and increased cell proliferation (Fig. 6). In summary, peritumoral lymphocyte aggregates could act as a lymphocyte reservoir, but could be unleashed by combining fibrolytic drugs to immune-based therapies.

The comprehensive annotation data and algorithms are available to expand the textural and lymphocyte analyses to other datasets, cancer types or even to a pan-cancer study. In summary, this study highlights how computational analysis of routine H&E staining can help to detect sampling fallacies, quantify tumor-infiltrating and tumor-surrounding lymphocyte infiltration, and discover novel histopathological associations both in the entire tumor sample and in the tumor margin.

## ONLINE METHODS

### TCGA ccRCC patient cohort

We collected histology images from TCGA portal (https://portal.gdc.cancer.gov/) and clinical and processed transcriptome data from https://gdc.cancer.gov/about-data/publications/panimmune^20^ and https://gdc.cancer.gov/node/905/ (PanCanAtlas; Fig. 1a). Samples were digitized mainly with an imaging resolution of ∼0.25 mm/px (n=497), except for 18 samples scanned at ∼0.50 mm/px, which were excluded (Fig. 1b, Supplementary Table 1). A feature matrix of all available data was built, consisting of only numeric or binary features as rows, with missing data reported as NA. Categorical variables were transformed into binary factors.

The somatic mutation calls (SNVs and indels) were collected from the supplementary table S1 by Ricketts *et al*^21^. Briefly, the dataset was assembled using six different algorithms (MuTect, MuSE, Pindel, Somatic Sniper, VarScan2 and Radia) from four centers. Mutation calls were not available for all patients with transcriptome and clinical data.

TCGA ethical statement and informed consent is available at https://www.cancer.gov/about-nci/organization/ccg/research/structural-enomics/tcga/history/policies.

### Image annotation strategy

TCGA H&E slides have been stained at distinct participating clinical sites and scanned with varying resolution and digital scanner types into SVS format. To ensure algorithm generalizability, we included annotations from all clinical centers.

First, we determined the main texture classes to annotate. We prioritized classification reliability and therefore minimized the number of tissue classes as often tiles included multiple texture types and some histological patterns commonly co-occurred, for example smooth muscle, fibrous stroma, and blood vessels. In addition, some patterns were challenging to differentiate reliably from individual tiles without larger context information for example torn, adipose, and necrotic tissue. We ended up with the following texture classes: renal cancer (“cancer”; n=11,755 image tiles, 29.7% of all tiles); normal renal (“normal”; n=6,313, 16.0%); stromal (“stroma”; n=3,027, 7.7%) including smooth muscle, fibrous stroma and blood vessels; red blood cells (“blood”; n=544, 0.9%); empty background (“empty”; n=11,609, 29.4%); and other textures including necrotic, torn, and adipose tissue (“other”; n=6,210, 15.7%; Fig. 1c). We annotated in total 39,458 randomly-selected tiles sized 300 × 300 px located in the center of a larger 900 × 900 px image to improve the annotation accuracy.

To quantify lymphocytes, we annotated 256 × 256 px tiles (n=25,095) containing none or few lymphocytes as “Low” (n=20,092, 80.1%) and the rest as “High” (n=5,003, 19.9%; Fig. 1d). As areas of high lymphocyte density were substantially less common, we speeded up annotation by extracting regions of high lymphocyte aggregates and from regions of low lymphocyte infiltrate using the open-source software QuPath^22^ 0.2.0. We selected ∼20 digital TCGA samples originating from various clinical sites. All texture and lymphocyte images were evaluated twice to minimize annotation errors.

### Texture classification

For texture classification, we trained a multi-class CNN. We employed the deep residual network ResNet as it has been commonly-used in computer vision tasks^23^. Transfer learning is the process of repurposing parameters of a previous algorithm to optimize training on a new dataset^24^. Here, we adapted transfer learning by combining the ImageNet-pretrained ResNet-18 infrastructure with a fully-connected layer, a rectified linear activation function (ReLU) activation and a softmax layer for prediction. Training occurred at all CNN layers, with the Adam optimizer tuned with a fixed learning rate of 10^−4^, batch size 4, and the cross-entropy loss function until the validation loss did not decrease for 5 consecutive epochs. We randomly cropped 256 × 256 px tiles from the annotation images and augmented these with horizontal-vertical rotation and without balancing texture classes. Models were composed with Python 3.9.1. with libraries Pytorch 1.9, Torch 1.11.0 and Torchvision 0.12.0.

The classification resulted in tessellated texture areas (Extended Data Fig. 13). For instance, cancer regions were disrupted by sporadic tiles of other textures. To smooth texture masks, we slid the 3 × 3 tiles’ window size and stride of 2 over the texture map and unified the texture class in each window by the most common texture. In some occasions, two tissue textures occurred equally often for instance 4 cancer, 4 stroma and 1 blood tiles. If the most common class was a tie between cancer and another texture, the cancer class was prioritized. If the most common class was a tie between stroma and another texture, the stroma class was prioritized except if the other was cancer. Stroma and cancer textures were prioritized as these occurred most in tiles of multiple textures. In other cases of tie, the pooled texture type was randomly selected from the equally most occurring textures.

### Lymphocyte classification

For the lymphocyte classification, we trained a binary-class CNN using the same model infrastructure and hyperparameters as was used in the texture classification. However, to quantify the lymphocyte infiltration in a continuous range [0-1], we used the argmax function on the sigmoid layer. Therefore, no post-pooling of lymphocyte masks was performed.

### Tumor margin

To analyze the histological and lymphocyte content immediately exterior to the tumor, we defined the tumor margin as the first two non-cancer tiles around each cancer tile with the maximum_filter function of the Python Scipy 1.8.1. library. The margin was 512 px or ∼128 µm wide. For reference, the average lymphocyte diameter is ∼10 µm. The remaining tissue not included as tumor or tumor margin was classified as the non-margin tissue. To avoid sampling bias, we included only samples with ≥1% normal texture.

### Model metrics

We divided annotation datasets into training (70%), validation (20%), and test (10%) sets. The final model fitness was evaluated in the test set by comparing classification accuracy and a confusion matrix.

### Statistical analysis

Median and interquartile ranges (IQR) were used to report average values and ranges. We compared two continuous variables with the Wilcoxon rank-sum test (unpaired, two-tailed) and three or more continuous variables with the Kruskal-Wallis test. We compared categorical variables with the χ^2^ test. To adjust p values, we used Benjamini–Hochberg correction. We evaluated survival with the Kaplan–Meier analysis (log-rank test). We limited comparisons with transcriptomic data to the protein-coding genes with a median expression value of >8 CPM (n=9,645). For pathway analyses, we included the Chromosome, Hallmark, PID, Reactome, Biocarta and KEGG gene sets v.6.2. We performed statistical analyses with R 3.5.1.

### Total lymphocyte infiltration normalization

The texture-specific lymphocyte infiltration varied by clinical centers partly due to differing texture proportions reflecting sampling conventions (Extended Data Fig. 7d). We harmonized total lymphocyte infiltration in two steps (Extended Data Fig. 7e). First, we multiplied the proportion of each texture area by its relative lymphocyte proportion. The relative lymphocyte proportion reflects the lymphocyte enrichment to each texture in comparison to other textures. Thus, the sum of relative lymphocyte proportion in blood + cancer, + normal + blood + other = 100%. In the second step, we divided the sum of the normalized lymphocyte proportions by the total sample area e.g.

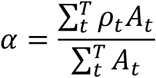

where *ρ* represents the relative lymphocyte proportion (%) and Аthe area (px) of the texture *t*, which is part of the texture list *T* ∈ [*blood, cancer, normal, stroma, other*].

To fully normalize staining differences, we categorized samples into “High” and “Low” infiltration based on their lymphocyte density compared to the clinical center median density. Small batch size increases the high risk for nonparametric data distribution. Therefore, we included only centers with more than 20 samples to the lymphocyte analyses.

## CODE AND DATA AVAILABILITY

The annotated texture and lymphocyte image data and algorithm parameters have been deposited to Zenodo https://zenodo.org/deposit/6528599. The code to run the CNN classifiers are available at https://github.com/vahvero/RCC_textures_and_lymphocytes_publication_image_analysis. The code and processed data to reproduce data analyses and figures are available at https://github.com/obruck/RCC_textures_and_lymphocytes_publication_data_analysis. The TissUUmaps visualization platform is available at http://hruh-20.it.helsinki.fi/rcc_texture_lymphocytes/.

## ACKNOWLEDGMENTS

We are grateful to the members of the Hematoscope Lab and Hematology Research Unit Helsinki for discussions and technical help.

## AUTHORS CONTRIBUTIONS

Conception and design: Ot.B., Os.B., S.M. Collection and assembly of data: Ot.B., Os.B., P.P. Image analysis: Ot.B., Os.B. Data analysis: Os.B. Visualization: Ot.B., Os.B. Manuscript writing: Os.B. Manuscript editing: All authors. Data interpretation: All authors. Final approval of manuscript: All authors.

