## Extended data for "Integrative Analysis of Histological Textures and Lymphocyte Infiltration in Renal Cell Carcinoma using Deep Learning"

Supplementary table 1. Samples excluded from analyses.

| TCGAid | errortype |
|---|---|
| TCGA-B8-5158 | low resolution |
| TCGA-B8-5159 | low resolution |
| TCGA-B8-5163 | low resolution |
| TCGA-B8-5164 | low resolution |
| TCGA-B8-5165 | low resolution |
| TCGA-BP-4770 | low resolution |
| TCGA-CZ-4853 | low resolution |
| TCGA-CZ-4854 | low resolution |
| TCGA-CZ-4856 | low resolution |
| TCGA-CZ-4857 | low resolution |
| TCGA-CZ-4858 | low resolution |
| TCGA-CZ-4859 | low resolution |
| TCGA-CZ-4861 | low resolution |
| TCGA-CZ-4862 | low resolution |
| TCGA-CZ-4863 | low resolution |
| TCGA-CZ-4864 | low resolution |
| TCGA-CZ-4865 | low resolution |
| TCGA-CZ-4866 | low resolution |
| TCGA-AK-3456 | necrosis |
| TCGA-BP-4804 | necrosis |
| TCGA-BP-4167 | necrosis&poor quality |
| TCGA-CJ-5671 | necrosis&poor quality |
| TCGA-A3-A8OV | no cancer |
| TCGA-A3-A8OX | no cancer |
| TCGA-B8-4619 | no cancer |
| TCGA-GK-A6C7 | no cancer |
| TCGA-A3-3328 | not ccrcc |
| TCGA-AK-3427 | not ccrcc |
| TCGA-AK-3427 | not ccrcc |
| TCGA-AK-3433 | not ccrcc |
| TCGA-AK-3440 | not ccrcc |
| TCGA-AK-3443 | not ccrcc |
| TCGA-AK-3447 | not ccrcc |
| TCGA-AK-3453 | not ccrcc |
| TCGA-AK-3465 | not ccrcc |
| TCGA-B0-4699 | not ccrcc |
| TCGA-B0-4834 | not ccrcc |

| | |
|---|---|
| TCGA-B0-5083 | not ccrcc |
| TCGA-B0-5117 | not ccrcc |
| TCGA-B2-3923 | not ccrcc |
| TCGA-BP-4334 | not ccrcc |
| TCGA-BP-4994 | not ccrcc |
| TCGA-AS-3777 | not typical ccrcc |
| TCGA-B2-4101 | not typical ccrcc |
| TCGA-B8-A54E | not typical ccrcc |
| TCGA-T7-A92I | not typical ccrcc + weird stain |
| TCGA-T7-A92I | not typical ccrcc + weird stain |
| TCGA-T7-A92I | not typical ccrcc + weird stain |
| TCGA-B0-5120 | poor quality |
| TCGA-BP-5201 | poor quality |
| TCGA-CJ-4891 | poor quality |
| TCGA-CJ-4892 | poor quality |
| TCGA-CJ-4893 | poor quality |
| TCGA-CJ-4899 | poor quality |
| TCGA-CJ-5672 | poor quality |
| TCGA-CJ-5675 | poor quality |
| TCGA-CJ-5676 | poor quality |
| TCGA-CJ-5677 | poor quality |
| TCGA-CJ-5678 | poor quality |
| TCGA-CJ-5679 | poor quality |
| TCGA-CJ-5680 | poor quality |
| TCGA-CJ-5681 | poor quality |
| TCGA-CJ-5682 | poor quality |
| TCGA-CJ-5683 | poor quality |
| TCGA-CJ-5684 | poor quality |
| TCGA-CJ-5686 | poor quality |
| TCGA-CJ-5689 | poor quality |
| TCGA-CZ-5452 | poor quality |
| TCGA-BP-4782 | poor quality, low proportion |

Supplementary table 2. Association of genomic alterations and blood texture in samples with normal tissue.

| genes | pvalue | Class 1 median | Class 2 median | Class 1 |
|---|---|---|---|---|
| mtor | 0.012 | 1.16 | 3.33 | Mutated |
| mutation_total | 0.12 | 2.65 | 3.77 | High burden |
| bap1 | 0.15 | 1.54 | 3.54 | Mutated |
| kdm5c | 0.16 | 6.87 | 3.16 | Mutated |
| vhl | 0.18 | 4.2 | 2.74 | Mutated |
| smarcb1 | 0.23 | 0.52 | 3.30 | Mutated |
| fat1 | 0.27 | 2.80 | 3.30 | Mutated |
| pik3ca | 0.29 | 2.05 | 3.31 | Mutated |
| nf2 | 0.30 | 2.17 | 3.30 | Mutated |
| kdm6a | 0.30 | 7.03 | 3.29 | Mutated |
| pbrm1 | 0.45 | 3.08 | 3.29 | Mutated |
| tp53 | 0.54 | 7.29 | 3.29 | Mutated |
| setd2 | 0.63 | 2.51 | 3.31 | Mutated |
| nfe2l2 | 0.63 | 3.16 | 3.30 | Mutated |
| pten | 0.90 | 3.28 | 3.29 | Mutated |
| stag2 | 0.92 | 4.14 | 3.22 | Mutated |
| ploidy | 0.99 | 3.83 | 3.29 | Diploid |

Supplementary table 3. Association of genomic alterations and blood texture in samples without normal tissue.

| genes | pvalue | Class 1 median | Class 2 median | Class 1 |
|---|---|---|---|---|
| pik3ca | 0.097 | 13.03 | 1.92 | Mutated |
| mutations_total | 0.17 | 2.63 | 1.81 | High burden |
| nf2 | 0.18 | 0.13 | 2.02 | Mutated |
| kdm5c | 0.25 | 3.29 | 1.89 | Mutated |
| smarcb1 | 0.26 | 0.30 | 2.02 | Mutated |
| pbrm1 | 0.32 | 1.97 | 2.02 | Mutated |
| setd2 | 0.42 | 1.29 | 2.02 | Mutated |
| nfe2l2 | 0.46 | 4.25 | 1.92 | Mutated |
| mtor | 0.50 | 2.64 | 1.97 | Mutated |
| bap1 | 0.55 | 1.92 | 2.04 | Mutated |
| vhl | 0.71 | 2.01 | 2.15 | Mutated |
| tp53 | 0.88 | 2.03 | 2.01 | Mutated |
| pten | 0.88 | 3.39 | 1.97 | Mutated |
| stag2 | 0.90 | 3.70 | 2.01 | Mutated |
| fat1 | 0.91 | 3.14 | 2.01 | Mutated |
| ploidy | 0.98 | 1.71 | 1.91 | Diploid |

Supplementary table 4. Association of genomic alterations and stroma texture in samples with normal tissue.

| genes | pvalue | Class 1 median | Class 2 median | Class 1 |
|---|---|---|---|---|
| pbrm1 | 0.0017 | 13.47 | 18.48 | Mutated |
| ploidy | 0.010 | 19.21 | 13.81 | Diploid |
| mutations_total | 0.050 | 15.21 | 17.85 | High burden |
| pten | 0.13 | 21.07 | 15.59 | Mutated |
| mtor | 0.15 | 11.49 | 16.35 | Mutated |
| kdm6a | 0.22 | 13.22 | 16.29 | Mutated |
| bap1 | 0.29 | 13.07 | 16.42 | Mutated |
| stag2 | 0.30 | 20.51 | 15.62 | Mutated |
| vhl | 0.31 | 15.39 | 17.67 | Mutated |
| nf2 | 0.40 | 20.70 | 15.62 | Mutated |
| tp53 | 0.41 | 21.13 | 15.62 | Mutated |
| kdm5c | 0.48 | 18.86 | 15.62 | Mutated |
| pik3ca | 0.51 | 18.93 | 15.62 | Mutated |
| setd2 | 0.55 | 16.30 | 16.02 | Mutated |
| nfe2l2 | 0.55 | 16.45 | 15.82 | Mutated |
| fat1 | 0.56 | 10.41 | 16.15 | Mutated |
| smarcb1 | 0.78 | 13.83 | 16.15 | Mutated |

Supplementary table 5. Association of genomic alterations and stroma texture in samples without normal tissue.

| genes | pvalue | Class 1 median | Class 2 median | Class 1 |
|---|---|---|---|---|
| mutations_total | 0.031 | 6.87 | 10.78 | High burden |
| setd2 | 0.038 | 6.40 | 10.38 | Mutated |
| stag2 | 0.060 | 16.79 | 9.54 | Mutated |
| smarcb1 | 0.085 | 0.00 | 10.07 | Mutated |
| ploidy | 0.15 | 10.78 | 10.20 | Diploid |
| nf2 | 0.18 | 0.99 | 10.067 | Mutated |
| pten | 0.47 | 10.70 | 9.75 | Mutated |
| vhl | 0.52 | 8.98 | 10.31 | Mutated |
| nfe2l2 | 0.58 | 14.13 | 9.99 | Mutated |
| pik3ca | 0.72 | 10.13 | 9.99 | Mutated |
| mtor | 0.76 | 9.95 | 10.07 | Mutated |
| pbrm1 | 0.78 | 10.20 | 9.54 | Mutated |
| kdm5c | 0.89 | 8.09 | 10.07 | Mutated |
| fat1 | 0.92 | 8.23 | 9.99 | Mutated |
| tp53 | 0.99 | 11.68 | 9.95 | Mutated |
| bap1 | 1.00 | 12.66 | 9.97 | Mutated |

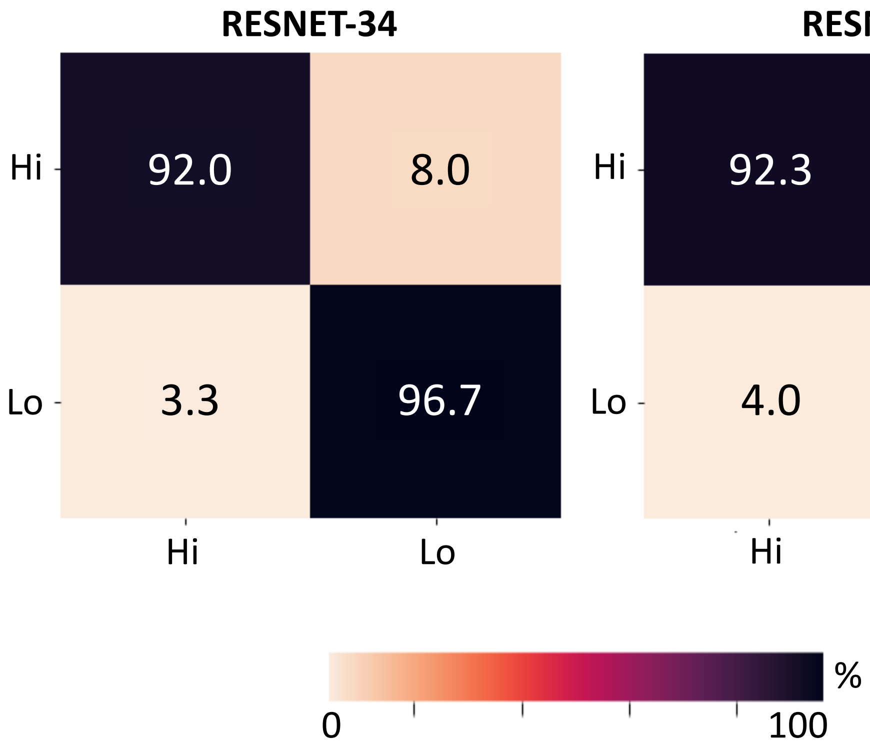


Extended Data Figure 1. Confusion matrix of the classification accuracy in the test set of the lymphocyte classifiers based on the ResNet-34 and ResNet-50 infrastructures

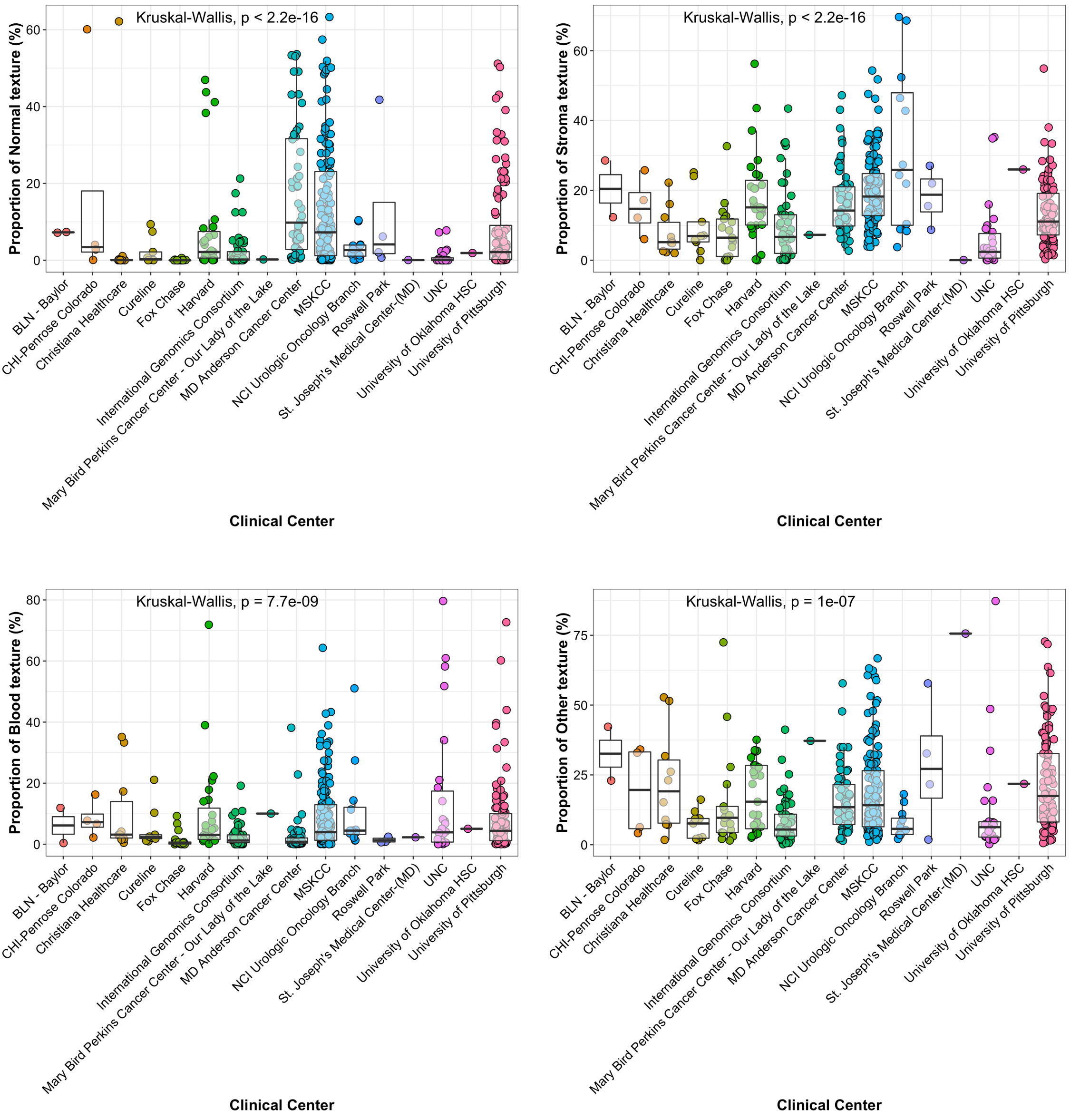


Extended Data Figure 2. Scatter and box plots to compare the proportion of normal renal tissue, stroma, blood and the other texture types by TCGA participating clinical sites. The box plots indicate the interquartile ranges and median values.

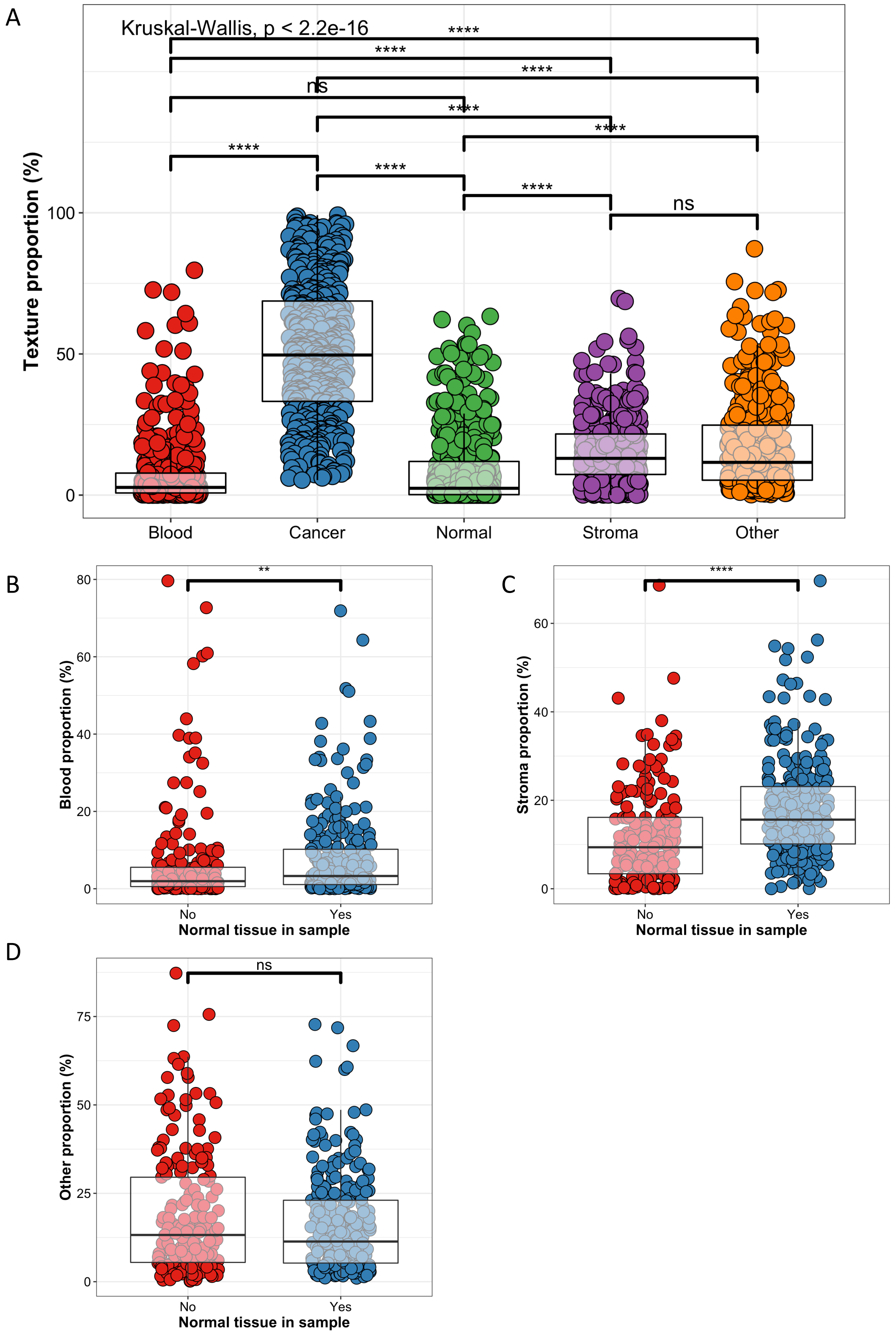


Extended Data Figure 3. (a) Scatter and box plots to compare the proportion of textures by tissue texture types. (b) Scatter and box plots to compare the proportion of blood, (c) stroma and (d) other texture types by samples without or with normal tissue. The box plots indicate the interquartile ranges and median values.

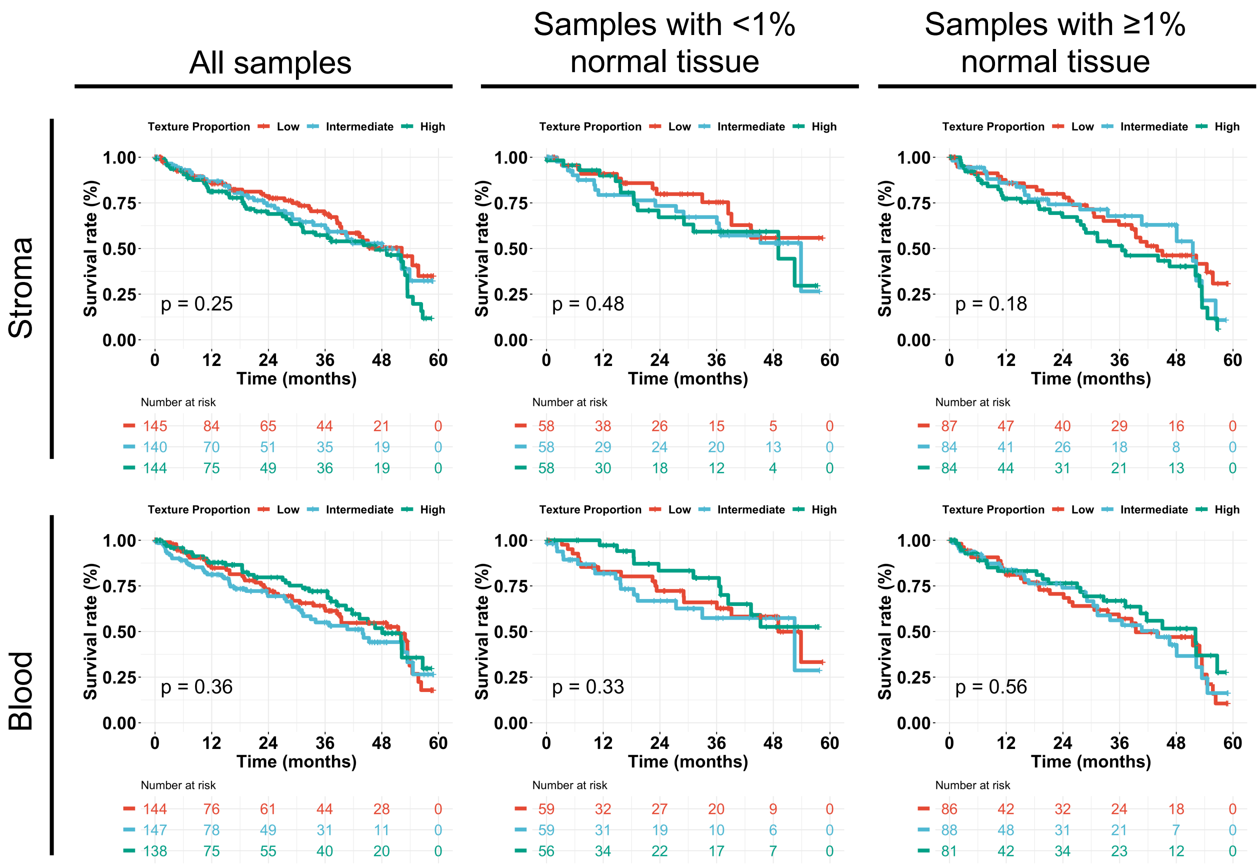


Extended Data Figure 4. Kaplan-Meier curves showing the survival association by the proportion of distinct tissue textures in all samples (left column), samples with <1% normal tissue (middle column) and samples with ≥1% normal tissue (right column).

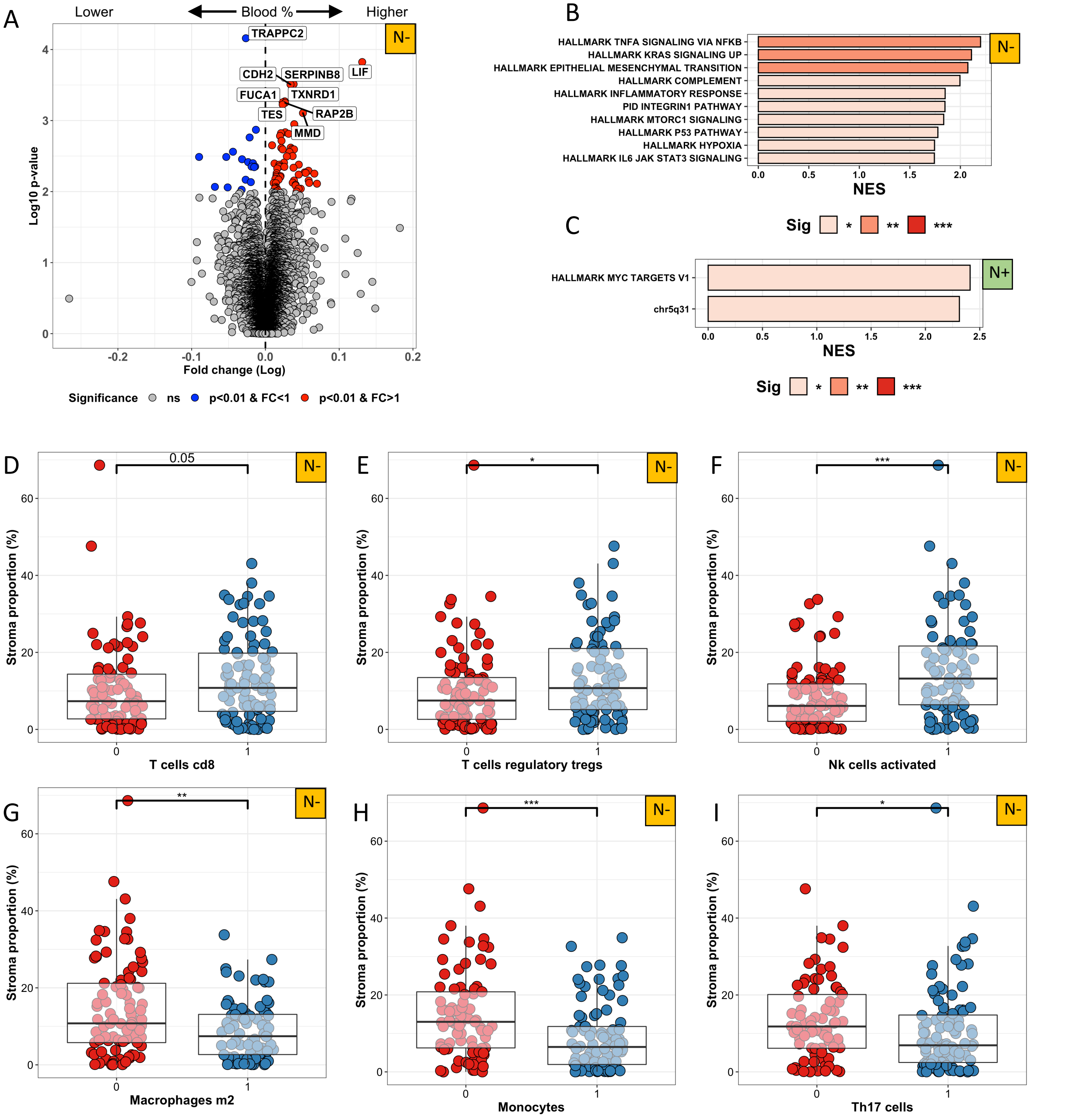


Extended Data Figure 5. (A) Volcano plot comparing the genes differentially-expressed in tumors with higher (right) to lower (left) than median hemorrhage proportion. Only samples without normal tissue are included. (B) Barplot indicating the normalized enrichment score (NES) of the gene pathways significantly associated with blood proportion in samples without and (C) with normal texture. (D) Scatter and box plots to compare the proportion of stroma texture by transcriptome-derived CD8+ T cell, (E) regulatory T cell, (F) activated NK cell, (G) M2-macrophage, (H) monocyte and (I) Th17 helper T cell proportion in samples without normal tissue.

Box plots: the lower and upper limit indicate the interquartile range and the center line the median value.

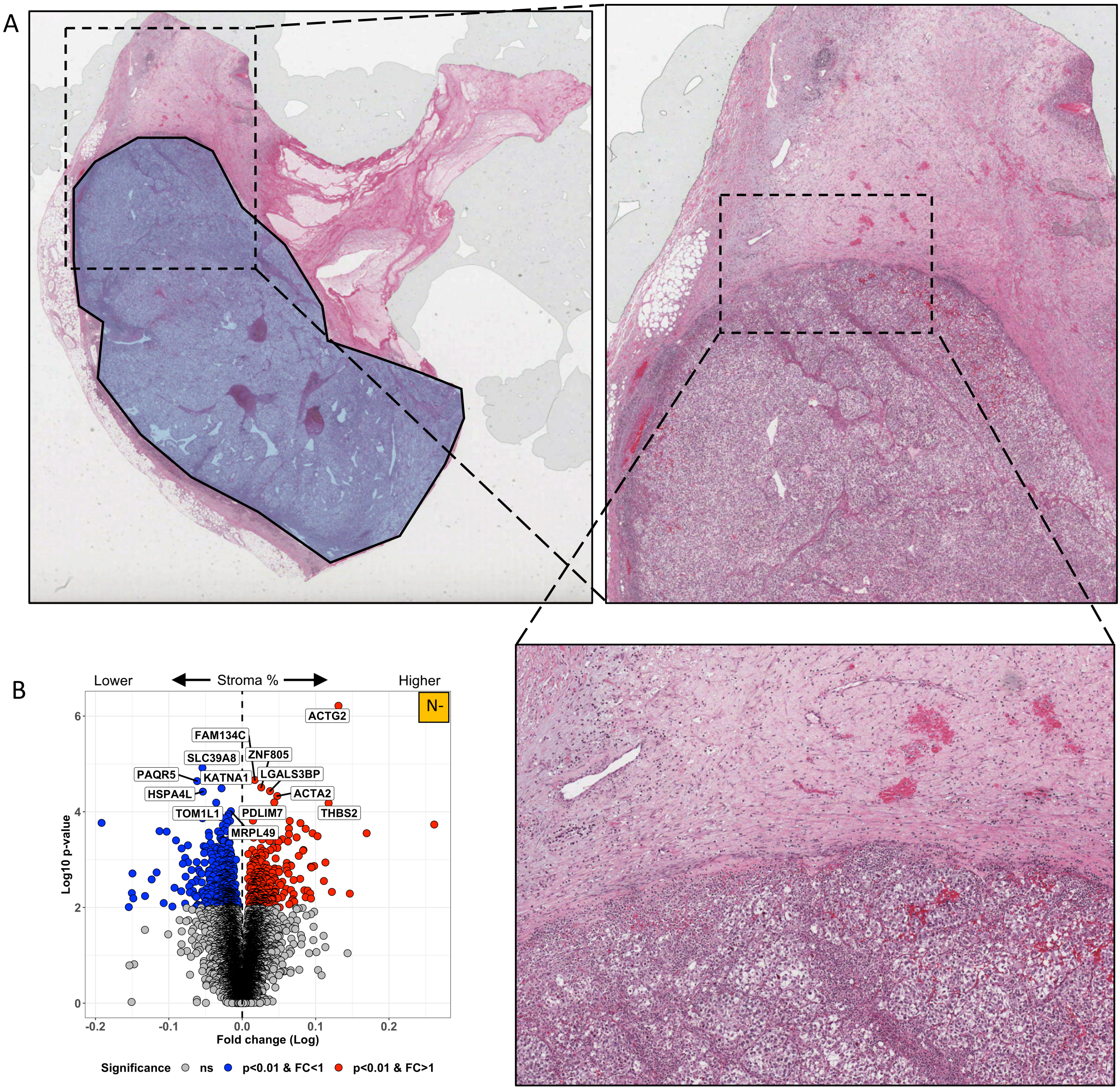


Extended Data Figure 6. (A) Example of the quantity and location of the intratumoral and peritumoral stroma from the H&E-stained clear-cell renal cell carcinoma TCGA-BP-4163. The blue mask represents the intratumoral tissue, while the rest represents the peritumoral tissue. (B) Volcano plot comparing the genes differentially-expressed in tumors with higher (right) to lower (left) than median stroma proportion. Only samples without normal tissue are included.

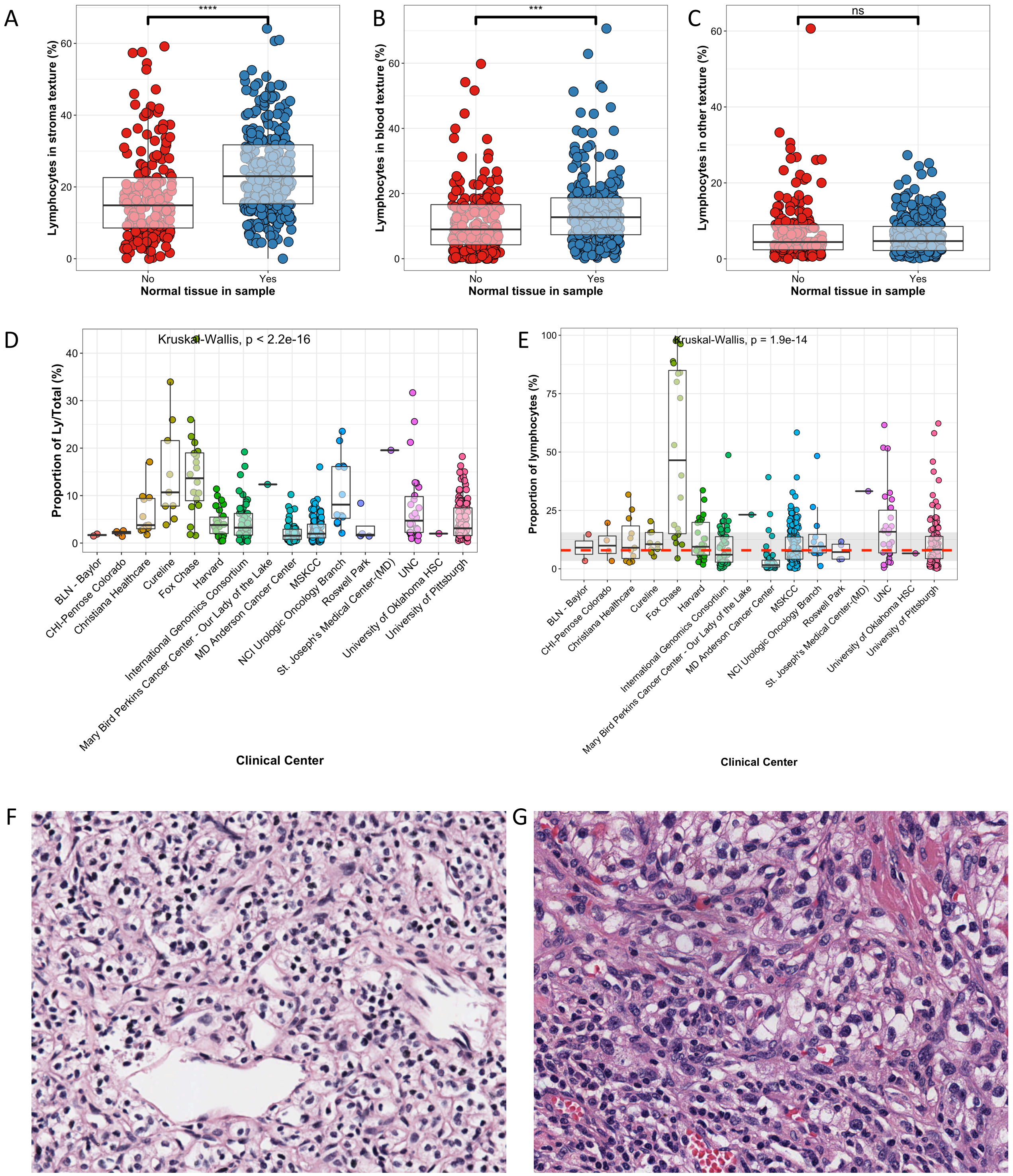


Extended Data Figure 7. The horizontal red dashed line indicate the median normalized lymphocyte proportion in all samples and the grey zone the 75% interquartile range. (A) Scatter and box plots to compare the proportion of lymphocytes in stroma, (B) blood, and (C) other texture by samples without or with normal tissue. (D) Scatter and box plots to compare the proportion of lymphocytes in the entire sample by TCGA participating clinical sites without and (E) with texture normalization. The same method was applied for all patients separately. The box plots indicate the interquartile ranges and median values. (F) Example H&E-staining of a nephrectomy section from the TCGA-AK-3434 patient from the Fox Chase center and (G) the TCGA-B0-5116 patient from the University of Pittsburgh.

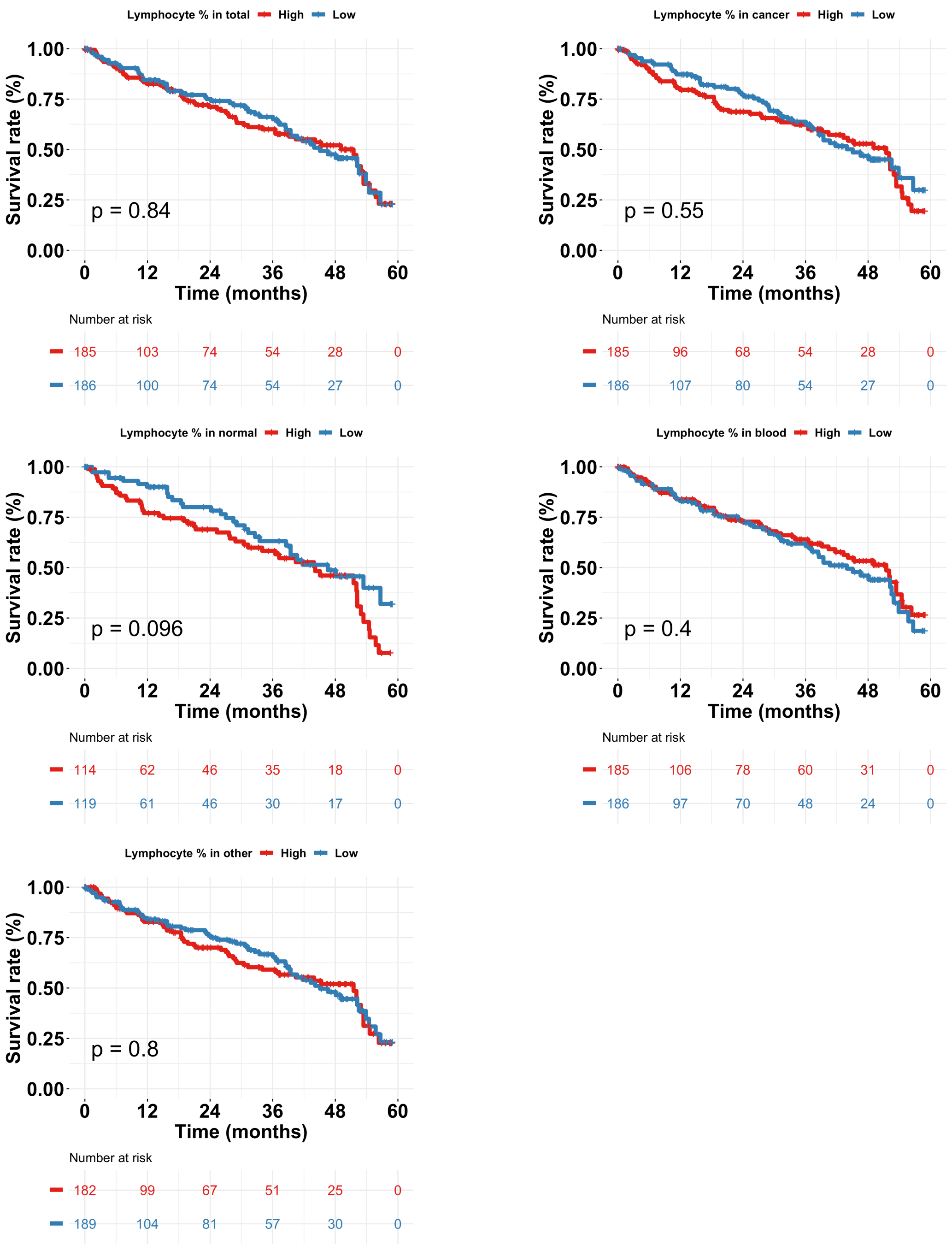


Extended Data Figure 8. Kaplan-Meier curves showing the survival association by the lymphocyte density in distinct tissue textures and in the entire sample.

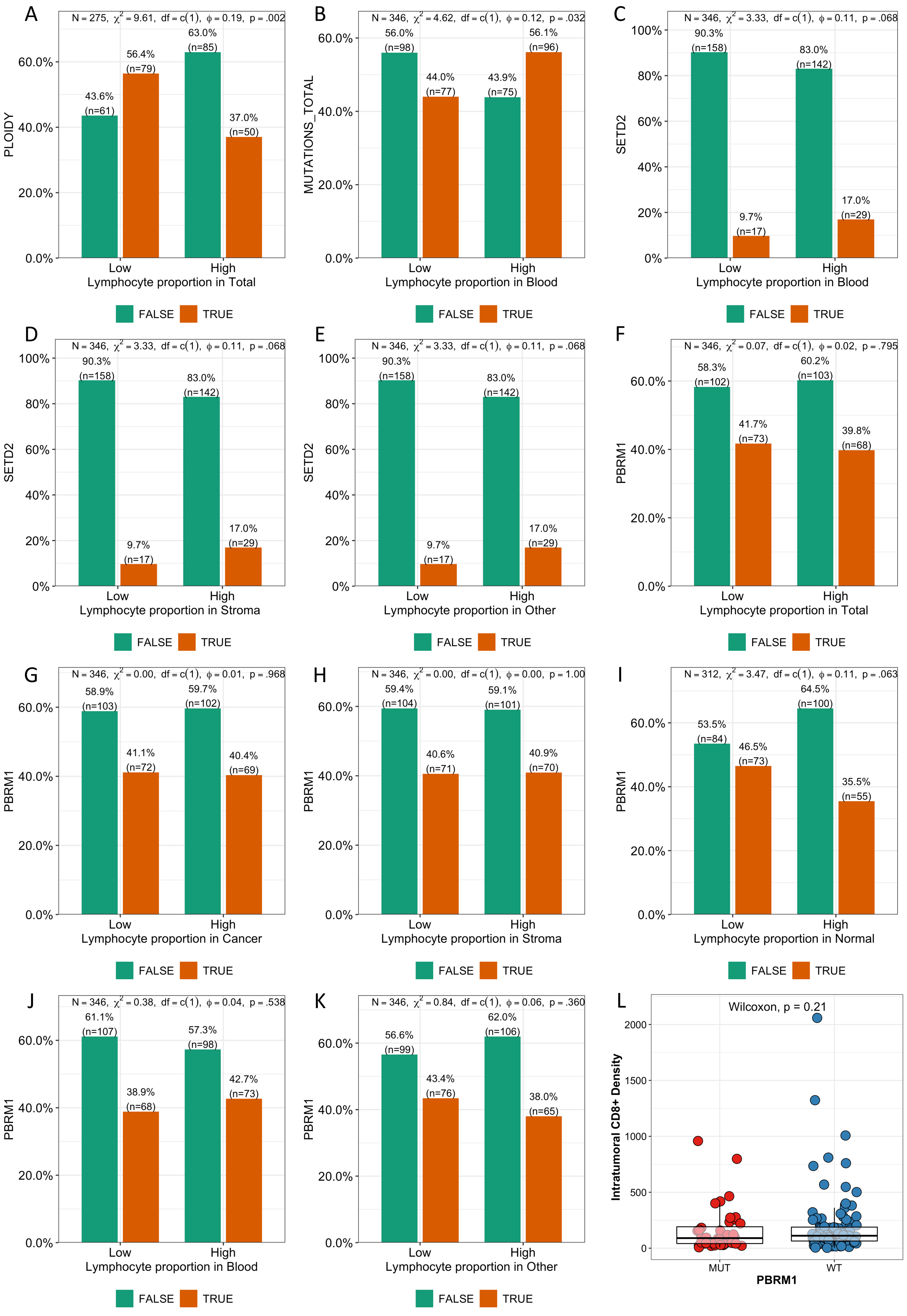


Extended Data Figure 9. (A-K) Barplots to compare lymphocyte density by textures and genomic variables. (L) Scatter and box plots to compare the proportion of intratumoral CD8+ T cell density by PBRM1 mutation status from the study by Braun *et al.*

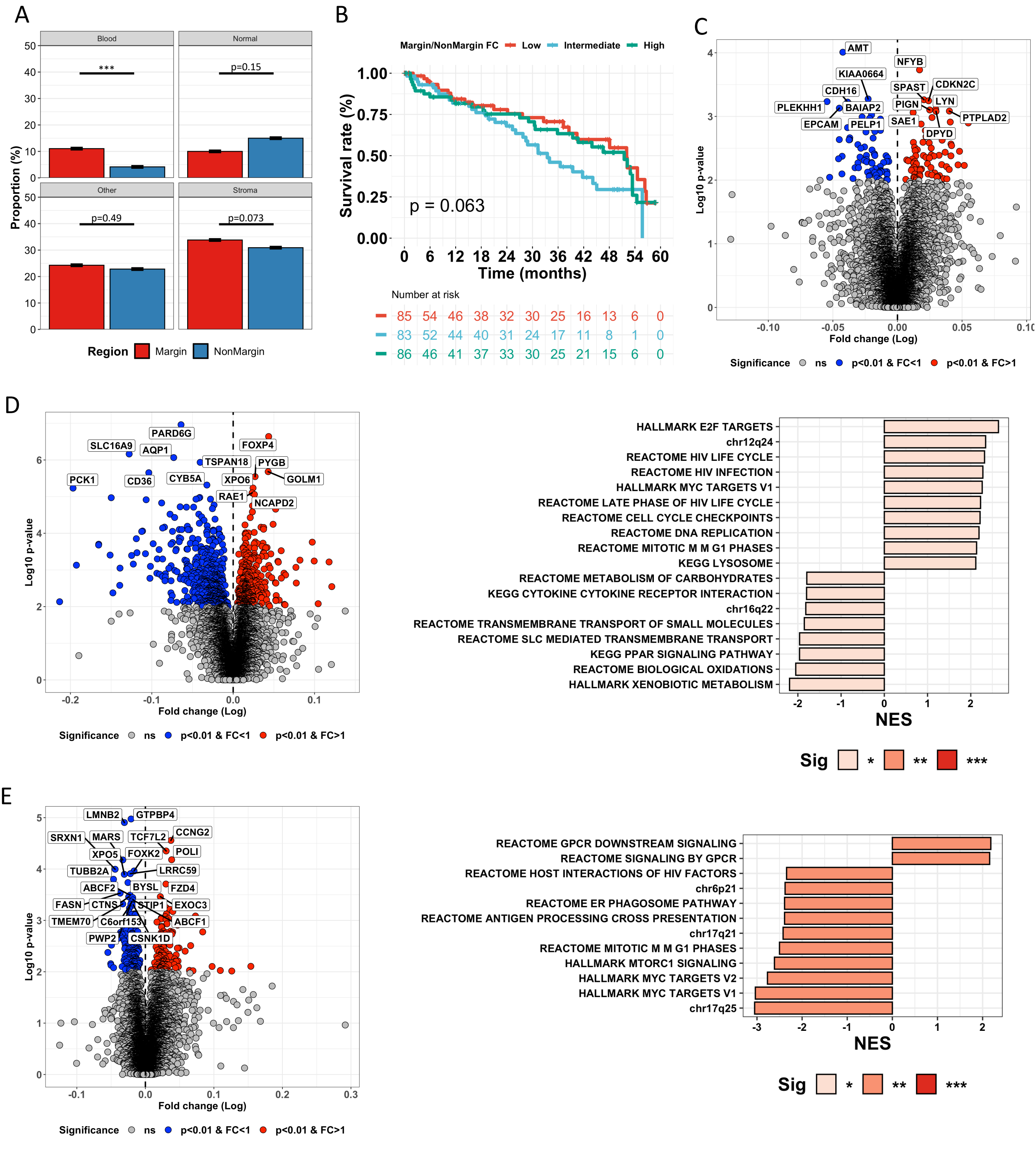


Extended Data Figure 10. (A) Box plots to compare the proportion of tissue textures in the tumor margin and non-margin. (B) Kaplan-Meier curves showing the survival association by the margin:non-margin ratio of the proportion of stroma tissue. (C) Volcano plot comparing the genes differentially-expressed in tumors with higher (right) vs. lower (left) margin sttroma proportion compared to non-margin. (D) Volcano plot comparing the genes differentially-expressed in tumors with higher (right) vs. lower (left) margin normal renal tissue proportion compared to non-margin. Barplot indicating the normalized enrichment score (NES) of the gene pathways significantly associated with the ratio of margin:non-margin normal renal tissue proportion. (E) Volcano plot comparing the genes differentially-expressed in tumors with higher (right) vs. lower (left) margin blood proportion compared to non-margin. Barplot indicating the normalized enrichment score (NES) of the gene pathways significantly associated with the ratio of margin:non-margin blood proportion.

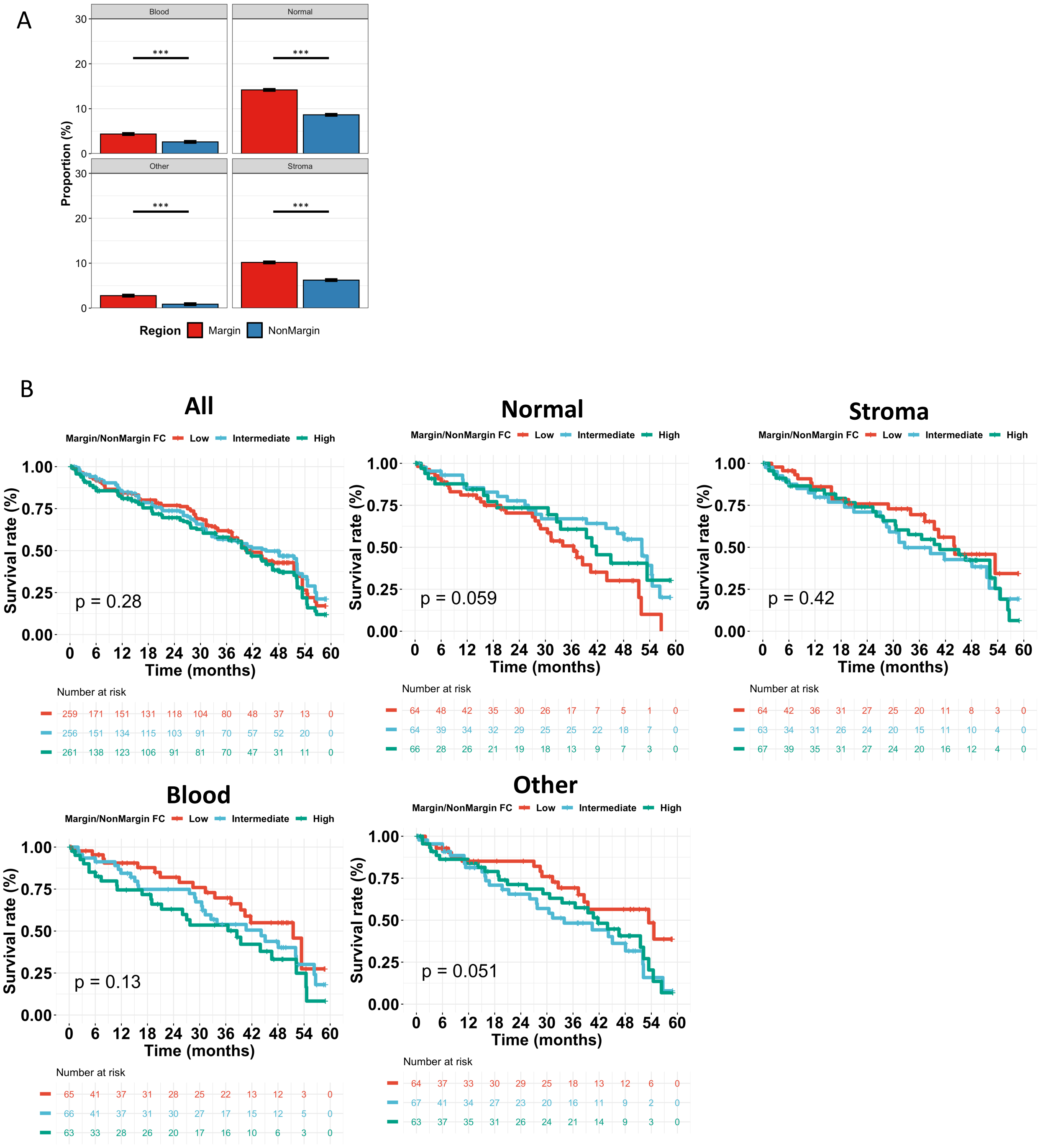


Extended Data Figure 11. (A) Box plots to compare the proportion of lymphocyte density in the tumor margin and non-margin. (B) Kaplan-Meier curves showing the survival association by the ratio of lymphocytes in the tumor margin compared to the non-margin.

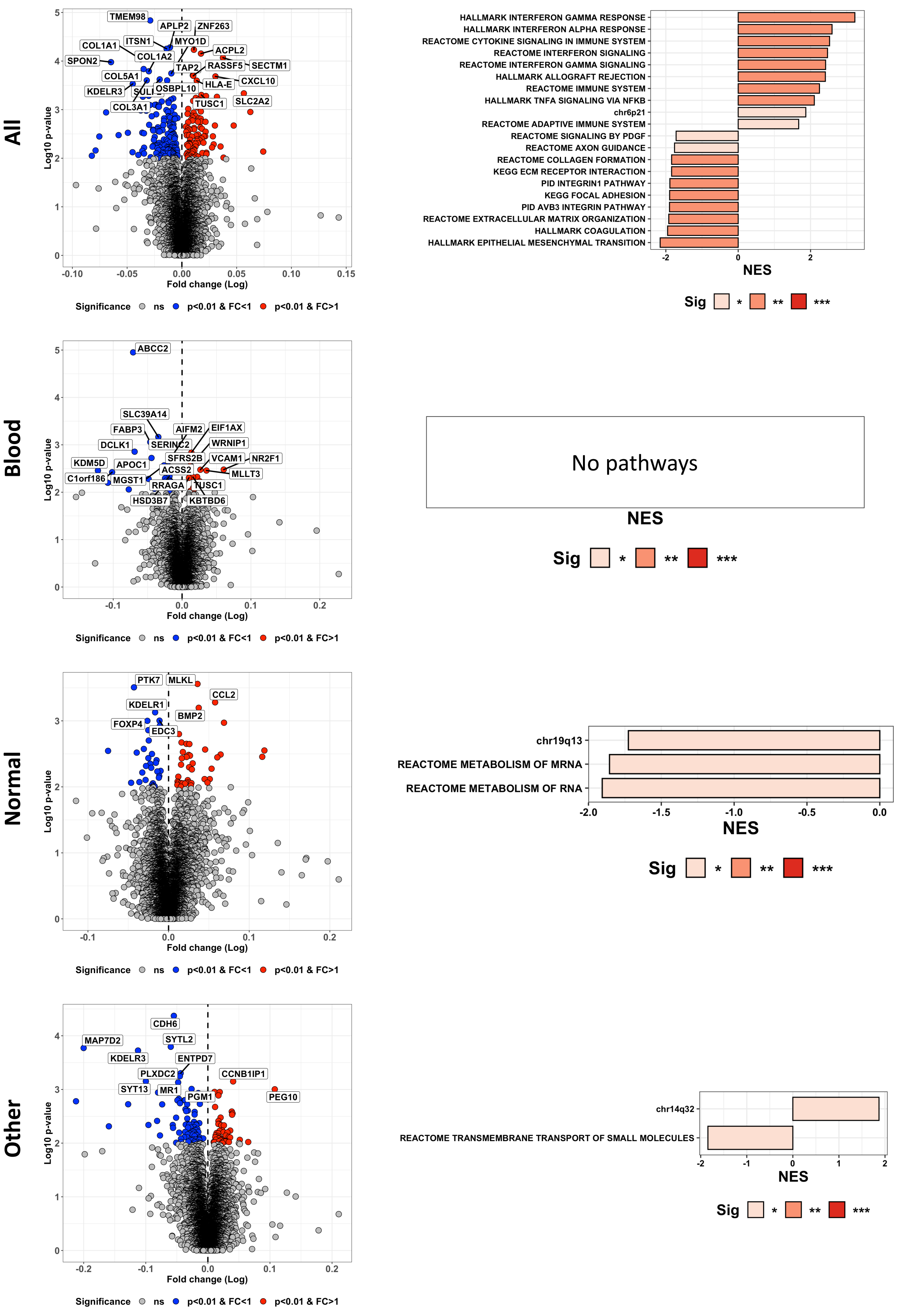


Extended Data Figure 12. Volcano plot comparing the genes differentially-expressed in tumors with higher (right) vs. lower (left) margin lymphocyte density compared to non-margin. Barplot indicating the normalized enrichment score (NES) of the gene pathways significantly associated with the ratio of margin:non-margin lymphocyte density.

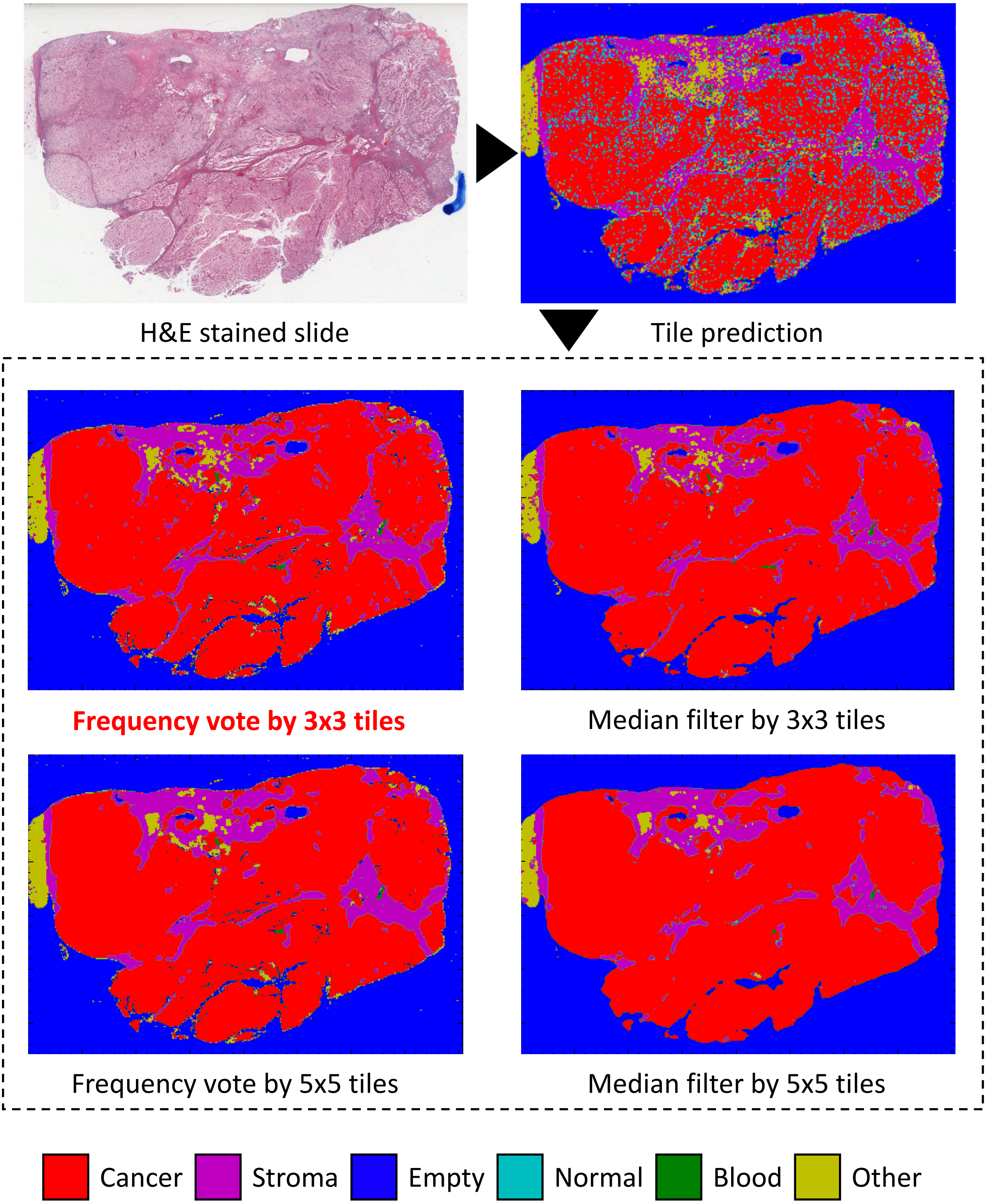


Extended Data Figure 13. Tissue textures are predicted for each tiles of the H&E-stained digital slide (top row). Many tiles display a mixture of multiple textures, which can be observed as a mosaic-like mask. The texture map has been averaged by selecting the most common texture ("frequency vote") or by selecting the median prediction value ("median filter") in combinations of 3x3 or 5x5 tiles. In this work, we employed the frequency vote by 3x3 tiles as it produced the best results based on visual review.
